# Functional network topography of the medial entorhinal cortex

**DOI:** 10.1101/2021.09.20.461016

**Authors:** Horst A. Obenhaus, Weijian Zong, R. Irene Jacobsen, Tobias Rose, Flavio Donato, Liangyi Chen, Heping Cheng, Tobias Bonhoeffer, May-Britt Moser, Edvard I. Moser

**Author notes:** **Corresponding authors** Horst A. Obenhaus,; Edvard I. Moser.

## Abstract

The medial entorhinal cortex (MEC) creates a map of local space, based on the firing patterns of grid, head direction (HD), border, and object-vector (OV) cells. How these cell types are organized anatomically is debated. In-depth analysis of this question requires collection of precise anatomical and activity data across large populations of neurons during unrestrained behavior, which neither electrophysiological nor previous imaging methods fully afford. Here we examined the topographic arrangement of spatially modulated neurons in MEC and adjacent parasubiculum using miniaturized, portable two-photon microscopes, which allow mice to roam freely in open fields. Grid cells exhibited low levels of co-occurrence with OV cells and clustered anatomically, while border, HD and OV cells tended to intermingle. These data suggest that grid-cell networks might be largely distinct from those of border, HD and OV cells and that grid cells exhibit strong coupling among themselves but weaker links to other cell types.

**Highlights:**

- Grid and object vector cells show low levels of regional co-occurrence
- Grid cells exhibit the strongest tendency to cluster among all spatial cell types
- Grid cells stay separate from border, head direction and object vector cells
- The territories of grid, head direction and border cells remain stable over weeks

## Introduction

The topographic organization of functional cell types in cortical areas has long been recognized as one of the fundamental organizational principles in the vertebrate brain (Penfield and Rasmussen, 1950) (for a review see Udin and Fawcett, 1988). In mammals, the large-scale, anatomical arrangement of function is most apparent in primary sensory cortices. These regions frequently possess an internal, large-scale topographic organization with respect to the stimulus space they cover, leading to the emergence of, for example, orientation maps, ocular dominance maps, and retinotopic maps in primary visual cortex (Hubel and Wiesel, 1965; Tootell et al., 1988) or tonotopic maps in primary auditory cortex (Hackett et al., 2011; Tsukano et al., 2017).

While these macro-scale patterns are relatively conserved across mammals, the extent to which single-cell tuning properties are topographically organized within each brain region varies from species to species (Jang et al., 2020; Scholl et al., 2013). For example, while the selective responses of neurons in the visual cortex to the orientation of edges follows a well-defined pinwheel-like organization in cats (Bonhoeffer and Grinvald, 1991; Ohki et al., 2006), similar patterns are generally thought to be absent in rodents (Van Hooser et al., 2005), although some, much looser, functional organization has recently been described in mice (Fahey et al., 2019; Ringach et al., 2016). The exact implications of differences in functional organization in these areas are still under debate, in particular how brain size, etiological and environmental demands on specific aspects of stimulus processing might favor one architecture over another (Ho et al., 2021; Koch et al., 2016; Swindale et al., 2000).

The underlying principles that dictate such patterning may derive from the constraints that speed and energy of information transfer impose on brain networks: Cell types that depend on frequent and dense information exchange may benefit from more direct anatomical proximity to each other than those that require less interaction (Luo, 2021; Song et al., 2014; Udvary et al., 2020; Wang and Clandinin, 2016). This is particularly true in brain regions in which circuit architecture might be less controlled by extrinsic factors than in many primary sensory cortices, and where the architecture instead follows more intrinsic organizational principles. In such higher-order regions, the study of functional topography can shape ideas about network topologies that determine both the local connectivity of cells as well as the emergence and maintenance of their tuning properties. One of the best illustrations of this concept in non-mammalian species is the discovery of a ring-shaped head-direction (HD) circuit in the centroid complex of fruit flies (Green et al., 2017; Kim et al., 2017; Seelig and Jayaraman, 2015), whose anatomical layout and wiring mirrors the predictions of ring attractor network models for directional tuning (Redish et al., 1996; Skaggs et al., 1995; Zhang, 1996). The anatomical layout of the fruit-fly head direction system minimizes wiring between similarly tuned dendrites.

In the mammalian brain, higher association cortices may follow similar principles. One such region where topographical organization may benefit computation is the mammalian medial entorhinal cortex (MEC), which contains a variety of distinct but functionally correlated cell types that together form a map of local space. These are grid cells (Hafting et al., 2005), whose firing fields tesselate the environment in equilateral triangles; border cells, which fire close to environmental boundaries (Solstad et al., 2008); object vector (OV) cells, which are active in fixed direction and distance from objects (Høydal et al., 2019); and HD cells that are active when the animal points its head towards specific directions (Sargolini et al., 2006; Taube et al., 1990). The abundance of these well-characterized cell types makes MEC an ideal candidate for the study of functional network topography, which in turn can lend evidence to the validity of network models describing the emergence of spatial coding in this region. Prominent models of HD and grid network topologies, so called continuous attractor network (CAN) models predict a dense, recurrent internal connectivity within networks of the same cell type, which in turn suggests that they should cluster anatomically (HD cells: Redish et al., 1996; Skaggs et al., 1995; Zhang, 1996; grid cells: Burak and Fiete, 2009; Couey et al., 2013; Fuhs and Touretzky, 2006; Guanella et al., 2007; McNaughton et al., 2006).

To elucidate these organizational principles, anatomically precise location and activity data is required. Two-photon (2P) *in vivo* calcium imaging is ideally suited for this because of its optical sectioning capability, high resolution, and its ability to record from hundreds of cells simultaneously over large field-of-views (FOVs) (Denk et al., 1990; Helmchen, 2009; Svoboda and Yasuda, 2006). Using large-scale 2P *in vivo* calcium imaging in head-fixed animals, recent studies have reported clustering of grid cells and associated tuning properties within MEC (Gu et al., 2018; Heys et al., 2014) supporting attractor network models that predict such patterns. However, due to the constraints imposed by the experimental setup, the analysis of other spatially modulated cell types was either impeded or rendered impossible (e.g., for HD cells) (Minderer et al., 2016). While macroscopic gradients have been described in freely-moving animals for both grid cells (Brun et al., 2008; Fyhn et al., 2004; Hafting et al., 2005; Stensola et al., 2012) and HD cells (Giocomo et al., 2014) a description of both the macro as well as micro scale anatomical organization of all known, spatially modulated cell classes in MEC and adjacent brain regions is still lacking.

Here we show results from imaging large areas of layers II/III of MEC and neighboring structures in mice using an upgraded version of the latest published miniaturized, portable two-photon microscopes (2P miniscopes) (Zong et al., 2017, 2021). This upgrade shares critical features with the most recent version of the miniscope introduced in the accompanying Resource article in this issue (Zong et al.). The modification to the previously published design enabled us to record from more than a hundred neurons simultaneously and resolve their spatial tuning properties at the same time as animals foraged freely in square open field arenas. The data enabled us to obtain a detailed record of anatomical organization within and between functional cell classes in the brain under quasi-natural conditions.

We acquired stable long-term, dual-channel recordings through chronic implants consisting of either a gradient refractive index (GRIN) lens combined with a micro prism, or a prism only (Low et al., 2014), in both cases positioned between MEC and the cerebellum. The second channel was used to visualize retrogradely labeled cells that project from MEC to the ipsilateral hippocampus (retrograde AAV, Tervo et al., 2016, carrying tdTomato, injected into the hippocampus). With these data we show that grid cells cover anatomically stable territories across superficial layers of MEC and neighboring regions that are distinct from those of other functional cell types, most notably object vector (OV) cells, which showed low co-occurrence with grid cells. Overall, all cell classes except for grid cells intermingle. Grid cells, but not other cell types, appear to cluster anatomically. The sharpest transitions between functional territories were visible at the border of MEC and neighboring parasubiculum (PAS).

These observations point to a unique topographical organization of grid and other spatial cell types in MEC. Grid cells seem to exist in relative isolation from other spatially modulated cells and may thereby form subnetworks that are relatively shielded from input of other cell classes.

## Results

### Large-FOV recordings with 2P miniscopes in freely moving mice

To obtain the anatomical and functional data required for topographical analyses of large numbers of spatially modulated cells in MEC, we used a fundamentally updated version of a previously published 2P miniscope (Zong et al., 2021), sharing decisive features with the even further developed version (“MINI2P”) introduced in the accompanying Resource article in this issue (Zong et al.) (Figure S1A and B). The present miniscope features a sufficiently large FOV for neural population recordings (width x height with GRIN+prism after motion correction (mean ± SD): 367 ± 40 × 558 ± 50 μm, Figure S1C) as well as sufficiently low weight (2.6 g without tether) to enable prolonged and unrestrained recordings in freely moving mice (Figure S1B). The weight decrease of this modified miniscope compared to its predecessors was achieved in both the scope itself, now made from a type of machinable plastic, as well as in the connection cable, which now contains a much thinner, tapered fiber bundle (“TFB”) with decreased overall diameter (for details see accompanying Resource article by Zong et al.). While lowering the weight, these design changes also drastically increased the degree to which the head-mounted assembly could be twisted and thereby permitted animals to exhibit more naturalistic foraging behaviors (see also accompanying Resource article). Some of the newest hardware features of MINI2P that are introduced in the accompanying Resource article, such as the even further enlarged FOV size, the built-in Z-focusing and the ability to shift FOVs systematically between sessions, were included in only a small subset of the present data (18 out of 212 recordings); importantly however, all present recordings had the lowered weight and the thinner, more flexible connection cable of MINI2P, allowing for good spatial coverage and reliable classification of spatially modulated cell types (see below).

### Chronic imaging of MEC and adjacent structures

Imaging was either performed in transgenic animals that expressed GCaMP6s ubiquitously in excitatory cells (Camk2a-tTA x tetO-G6s, “Transgenic”) or wild type mice that had been injected with recombinant AAVs expressing GCaMP6m (“Virus”, synapsin promoter) (Figure 1A). To gain access to the superficial layers of MEC and neighboring brain regions we chronically implanted optical relays consisting of a GRIN and prism doublet (diameter 1 mm, overall length ∼4.7 mm, Figure 1A bottom, Figure S1C right) or prisms only (1.3×1.3×1.6 mm) in between cerebellum and MEC, such that the prism surface was centered on and flush against the superficial layers of MEC (Figure 1 A to C). This approach enabled us to image MEC chronically for weeks and made it possible to capture the activity of over a hundred cells simultaneously after filtering for signal-to-noise ratio (SNR) (example in Figure 1D and E) (median SNR across 24,820 cells, virus (median): 5.6, transgenic: 4.8, Figure S1D; number of cells across n=133 sessions after filtering, virus (mean ± SD): 112 ± 68, transgenic: 161 ± 96, Figure S1E). To improve anatomical specificity *in vivo* we labelled superficial layer MEC cells selectively by injecting retrogradely transported AAVs carrying tdTomato into the ipsilateral hippocampus (Figure 1A, Figure S1F). This resulted in labeling of mostly layer III and some layer II cells in MEC (Figure 1C and Figure S1G) and thereby allowed us to distinguish MEC from neighboring structures like the medially located parasubiculum (PAS) as well as layer II from layer III by expression differences (i.e., abundance of tdTomato cell labeling in layer III compared to layer II).

**Figure 1.**
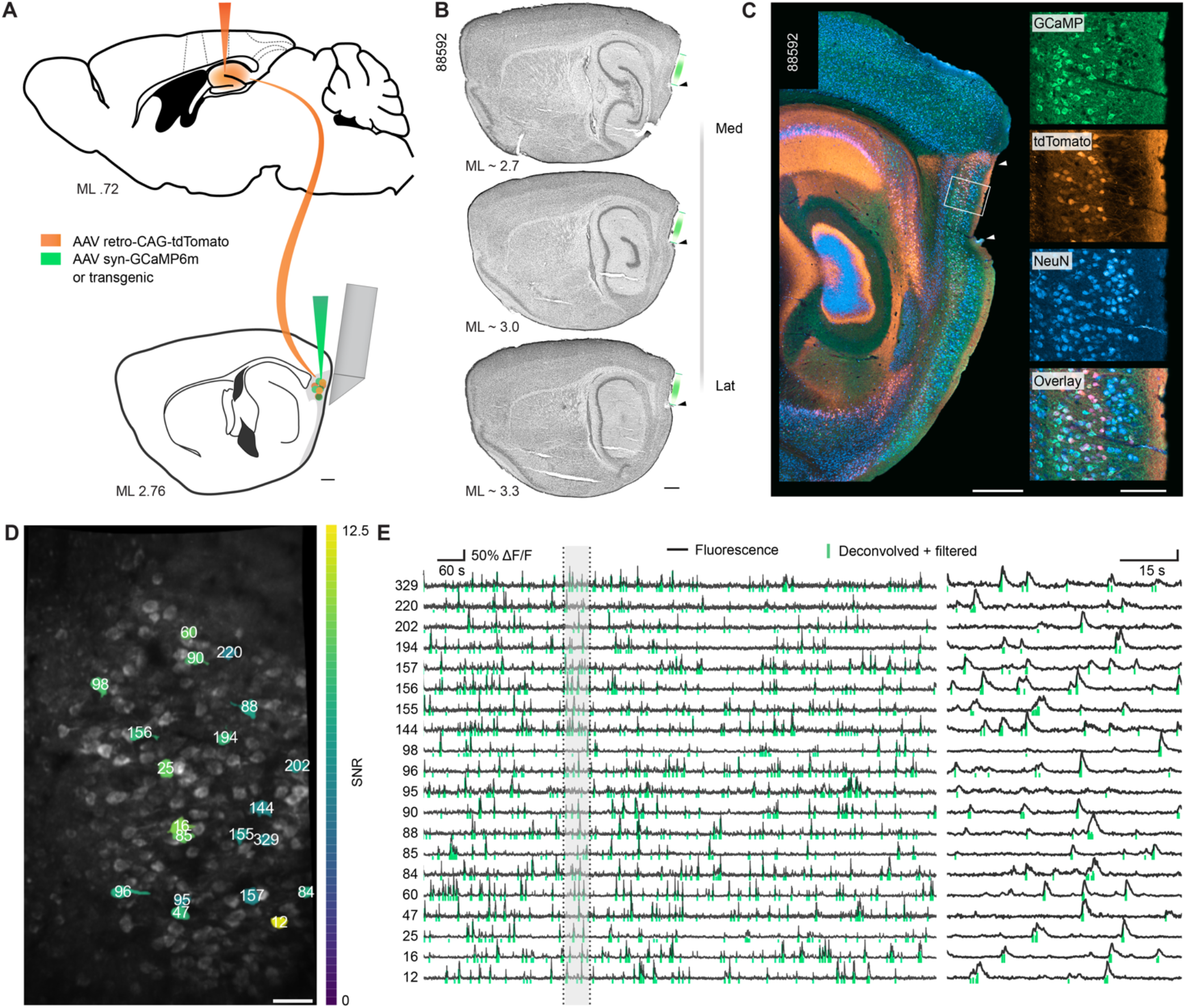
Calcium imaging with two-photon miniscope in medial entorhinal cortex (MEC). **(A)** Sagittal brain section schematics at two medio-lateral (ML) positions. Combined GRIN+prisms were chronically implanted between cerebellum and MEC, with the prism flush against superficial MEC. A GCaMP6m expressing virus, locally injected into superficial MEC, or transgenic mice expressing GCaMP6s were used (green). Retro-AAV expressing tdTomato was injected into the ipsilateral hippocampus to label hippocampus-projecting MEC neurons (orange). Scale bar 500 μm. **(B)** Nissl-stained, sagittal brain slices from one animal showing the impression of the implant against superficial MEC. Three sections are shown from medial (Med) to lateral (Lat). The approximate medio-lateral (ML) position is indicated at the bottom left of each section. Green lines indicate the maximum extent that could be imaged through the GRIN+prism configuration. Black arrow tips indicate the most ventral position of the prism. Mouse ID is shown in top left. Scale bar 500 μm. **(C)**Example immunofluorescence in a brain slice of a GCaMP6 transgenic mouse (same animal as in (B)). White arrow tips indicate the dorsal end of MEC (top) and ventral edge of the prism (bottom). Insets to the right show magnification of the white box. GCaMP (green), tdTomato (orange), NeuN (cyan) and overlay (GCaMP + tdTomato + NeuN) are shown separately. While GCaMP positive cells are found across several layers, tdTomato expression is mainly restricted to layer III. Scale bar 500 μm, insets: 125 μm. **(D)** Example maximum intensity projection of one open-field session with a total of 213 detected cells (after SNR filtering). The FOV is densely covered by GCaMP labeled cells. Colored regions show cells whose signal is shown in (E) with numbers on top of cell bodies indicating Suite2p assigned cell IDs. Scale bar 50 μm. **(E)** Fluorescent (normalized ΔF/F, black traces) and deconvolved and filtered signal (green vertical bars) of 20 example cells (colored ROIs in (D)). Total session length ∼1200 seconds. Shaded region (60 second excerpt) is magnified on the right.

### 2P miniscope imaging enables analysis of all spatial cell types

Spatially modulated cell types have classically been studied in animals that were allowed to engage in unrestrained foraging behavior in open field arenas. Due to the low weight of the 2P miniscope and its thin and flexible connection cable, we were likewise able to let animals run in large square boxes over tens of minutes (Figure 2A). Open field arenas were mostly 80×80 cm in size, although a few sessions were run in 60×60 and 100×100 cm arenas. Animals readily engaged in natural exploratory behaviors in all of these arenas while foraging for cookie crumbs (Figure 2A and B and Figure S2A). Optimal coverage was achieved after just ∼20 minutes (Figure 2B and C, 15 animals, 203 sessions), which greatly facilitated the interpretation of spatial modulation in recorded cells (for a detailed analysis of behavior and arena coverage see Figure 1 and S1 of the accompanying Resource article by Zong et al.). Average and maximum speed of animals during recordings matched that of previous reports using chronic implants in mice (for example chronic implantations of tetrode bundles for electrophysiological recordings, Dannenberg et al., 2019) (Figure S2B). Moreover, the behavior stayed stable over long durations such that we could record multiple sessions in a row (Figure S2C) and animals required only a few resting periods when inside the open field arena (Figure S2D). This was of pivotal importance for the analysis of object vector cells, which are analyzed by running consecutive sessions (usually one baseline session without objects and two object sessions) to follow the emergence of object-related firing fields and their re-allocation when the object(s) are moved between sessions (Høydal et al., 2019).

**Figure 2.**
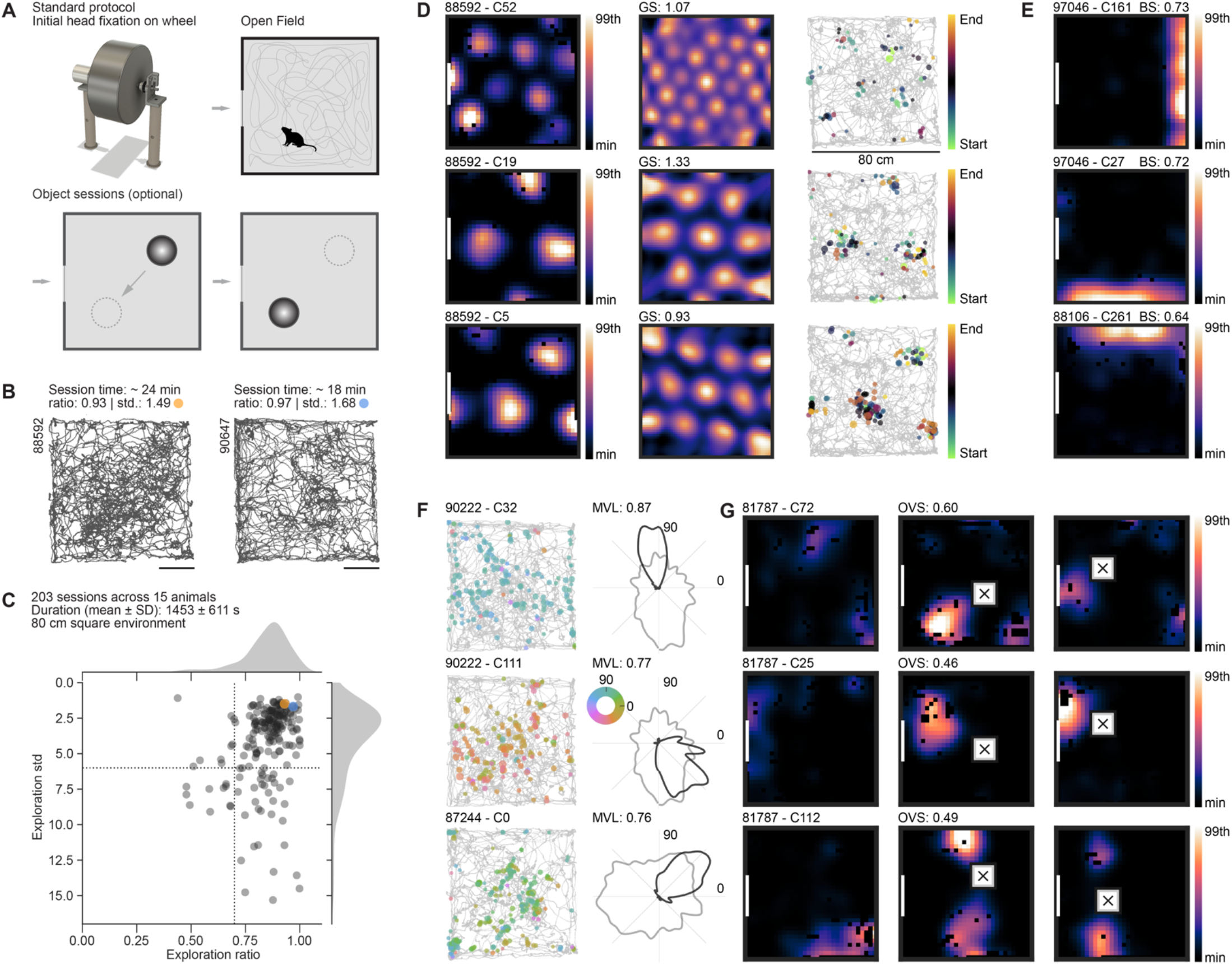
2P miniscope imaging of all functional cell types in MEC during unrestrained behavior. **(A)** Top: Schematic showing the usual order that recording sessions were run in. After initial head fixation on a wheel and attachment of the miniscope, animals were moved into a square open field arena (“Standard Protocol”), sometimes combined with further sessions with an added object (“Object sessions (optional)”), which was moved between sessions or a single session with two objects at a time to screen for object vector (OV) cells. **(B)** Example path plots of two recorded sessions in two different animals (animal number indicated in top left). Session time, exploration ratio (“ratio”, how many bins of all bins were visited) and exploration standard deviation (“std”, standard deviation of the time the animal spent across all bins) are indicated in the title. The colored dots (orange and blue) are referenced in (C). Scale bar 20 cm. **(C)** Analysis of open field coverage for 203 open field sessions across 15 recorded animals (80 cm square open field arena, duration 1453 s ± 610.6 (mean ± SD)). Plotted is the exploration ratio (“ratio”) versus exploration standard deviation (“std”). Animals chosen for this study exhibited on average good behavior, covering the open field homogeneously throughout the length of the session. Stippled lines show arbitrary cutoffs, splitting into quadrants of “better” (top right) and “worse” coverage (bottom left) (see also Figure S2A for further examples). The two sessions shown in (B) are indicated in orange and blue. **(D-G)** Example grid (D), border (E), head direction (HD, (F)) and object vector (OV, (G)) cells recorded during ∼ 20 min of free foraging in an 80×80cm open field. **(D)** Grid cells: spatial tuning maps (left column) for grid cells of a module with short grid spacing (top) and two co-recorded grid cells from a module with larger spacing (middle and bottom). Colormaps show spatial tuning maps (minimum to 99^th^ percentile). White lines indicate the position of a (single) cue card, fixed to the wall of the box. Middle column: 2D autocorrelations of spatial tuning maps on the left (“GS”: grid score). Right column: path and deconvolved signal (amplitude indicated by dot size, session time indicated by color). **(E)** Border cells: three example spatial tuning maps of border cells with border score (“BS”) indicated on top of the spatial tuning map. White line indicates the position of a cue card. **(F)**HD cells: path and deconvolved signal (left) and tuning curves (right) of three example head direction cells. Tuning strength (mean vector length, “MVL”) is displayed on top of the tuning curve. Deconvolved signal is shown color-coded by head direction on top of the path plot. Tuning curves show normalized occupancy (grey line) and the cell’s directional tuning curve (black line). **(G)** OV cells: spatial tuning maps of three object vector cells, two with one field (top and middle), and one with two fields (bottom), all maintaining vector relation to the object when the object is moved between the second and third trial. White line indicates position of a cue card; white square with black cross indicates position of the object. Object-vector score (“OVS”) is displayed above the middle tuning map. Colormaps have the same limits across all sessions for one cell (range of baseline session).

In this way we were able to analyze all major spatial classes, namely grid (Figure 2D), border (Figure 2E), HD (Figure 2F) and OV (Figure 2G) cells. To assign cells to either cell type (i.e., grid, HD, border or OV) we compared their spatial or directional tuning scores (e.g., grid score for grid cells) to shuffled distributions created by shifting the tracking and fluorescence signal for every cell against each other in random time intervals (see *STAR* methods). For grid, border, and HD cells we used single criteria with 95^th^ percentile shuffling cutoffs, if not otherwise indicated, and combined criteria were used to classify OV cells (see *STAR* methods). Scores matched that of previous reports using electrophysiological methods (Rowland et al., 2018). Due to the high SNR of the recorded signal and precision increases achieved by deconvolving the fluorescent traces, we were able to resolve closely spaced firing fields of grid cells (see example in Figure 2D, top row) as well as precise head direction tuning (see example in Figure 2F, top row), despite the slow time courses of calcium dynamics (for further examples see also Figure 5 and Figure S5 in the accompanying Resource article by Zong et al.). Cells that were recorded across adjacent recording sessions exhibited reproducible, stable tuning (measured both for head direction modulation as well as spatial tuning map correlation, Figure S2E and F). Due to the high throughput achievable in our large-scale imaging approach we were able to determine the properties of tens of object vector cells (OVCs) simultaneously while obtaining field-object distance and angle distributions that closely resemble that of previously published reports (Høydal et al., 2019) (Figure S2G).

Our imaging approach thus enabled high throughput identification of spatial cell types and anatomical position of over a hundred simultaneously recorded cells in MEC and adjacent structures.

### Object vector cells and grid cells segregate

While imaging in MEC and neighboring regions we noticed that recordings that showed an abundance of grid cells in one FOV (∼20% of all imaged cells) often would have comparably few OV cells (example in Figure 3A, 98 grid cells versus 3 OV cells) and, conversely, recordings with multiple OV cells would have only few co-recorded grid cells (example in Figure 3B, 27 OV cells versus 9 grid cells). Across all recordings this divergence was most visible in animals in which 10% or more of either cell type were recorded on average (Figure S2H and Figure 3C). While differences in cell type composition were visible between other pairs of spatially modulated cell classes as well (Figure 3D and Figure S3A and B), the most striking difference in cell numbers was found for grid versus OV cells (Figure 3C and D). This difference was maintained when we included conjunctive cells (i.e., those that crossed cutoff criteria for more than one cell class) (Figure S3B).

**Figure 3.**
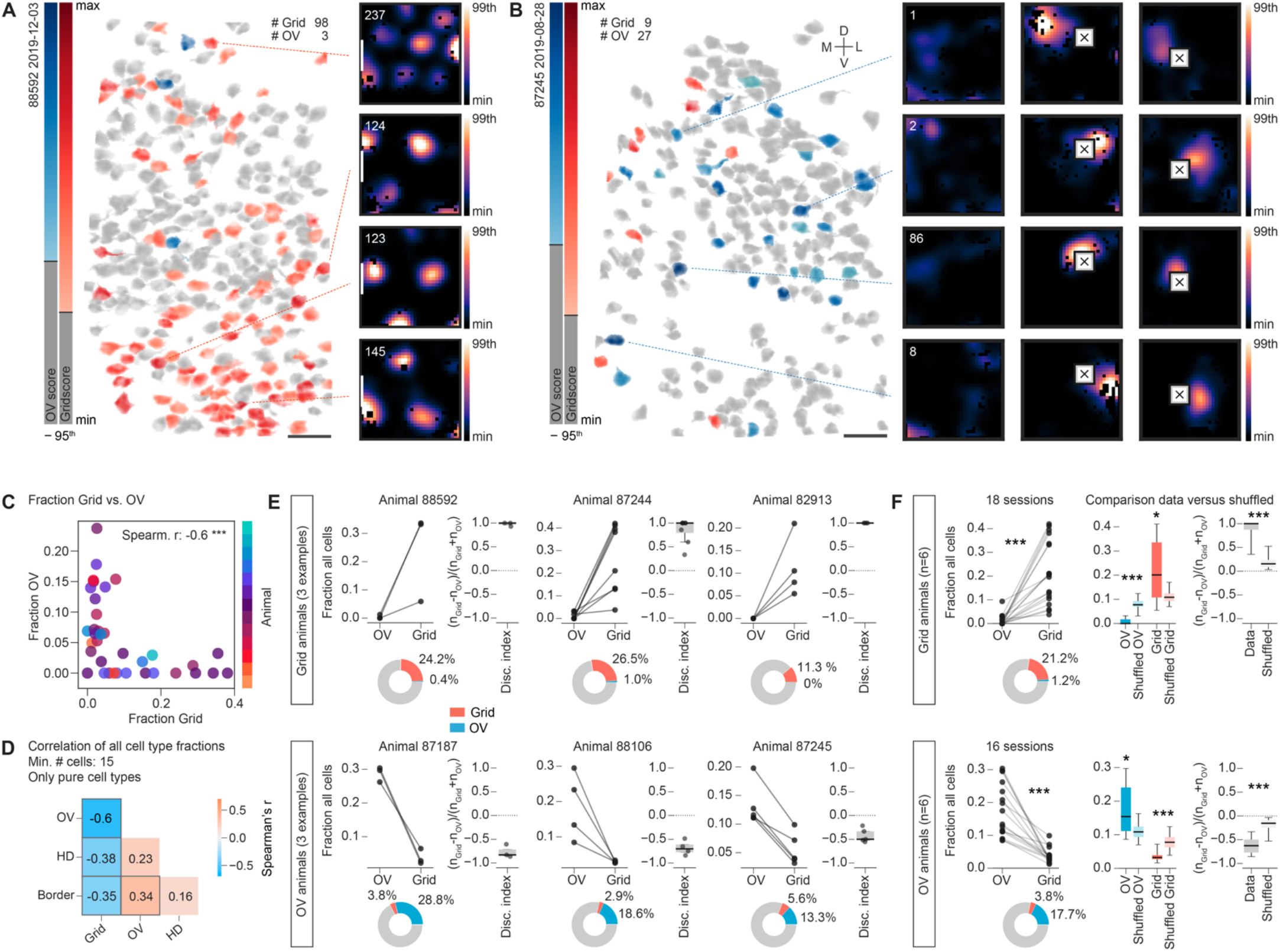
Grid cells and object vector (OV) cells occupy discrete regions of MEC. **(A+B)** Anatomical distribution of grid and OV cells in two different animals (animal name and recording date at top left) and example spatial tuning maps of cells in each session. Cells are color coded by tuning strength (OV score (blue), grid score (red), transition from grey to colored area on color bars shows 95^th^ percentile shuffling cutoff). Grey colored cells did not meet any of the cutoff criteria for classification as grid or object vector cells. **(A)** Recording with multiple grid cells and few OV cells. Numbers on top indicate number (#) of detected grid and OV cells. Spatial tuning maps towards the right in each panel show examples of four grid cells with ROI number indicated in the top left corner (scaling / display for spatial tuning maps as in Figure 2). Scale bar for cell map 50 μm. **(B)** As in (A), but for a recording in which multiple OV cells, but few grid cells, were identified. Spatial tuning maps towards the right in each panel show examples of four OV cells. Scale bar for cell map 50 μm. **(C)** Fraction of (pure) grid and object vector (OV) cells in 36 sessions (combined open field and object sessions) across 13 animals. Each dot represents a session and animals are coded by color. Spearman’s correlation is shown in top right (Spearman r=-0.596, n_Grid_=n_OV_=36, p=0.00013***). See also Figure S3A and B. **(D)** Summary of correlations as in (C), but for all cell types. Spearman’s r for each pair of comparisons is color coded and shown as text in the center of the box (filtered by MEC, minimum cutoff of 15 cells after filtering, only pure cell types). Thick grey border around some squares indicates significance (p<0.05) in Spearman’s correlation result (Grid vs. Border: r=-0.35, n_Grid_=n_Border_=124, p=5.47e-05***; Grid vs. HD: r=-0.38, n_Grid_=n_HD_=124, p=1.27e-05***; Grid vs. OV: r=-0.596, n_Grid_=n_OV_=36, p=0.00013***; OV vs. Border: r=0.34, n_OV_=n_Border_=36, p=0.046*; OV vs. HD: r=0.23, n_OV_=n_HD_=36, p=0.17*ns*; HD vs. Border: r=0.16, n_HD_=n_Border_=124, p=0.084 *ns*). See also Figure S3A and B. **(E)**Fraction of (pure) grid and OV cells and the discrimination index over grid and OV cells in each session across 6 example animals (minimum cutoff of 15 cells), with higher fractions of grid cells than OV cells (top row, 3 “Grid animals”) or the opposite (bottom row, 3 “OV animals”). A summary across all sessions and animals is shown in (F). See also Figure S3C for 6 additional animals. Pie charts underneath each line plot show the average percentage of grid and OV cells across sessions. Boxes in box plots extend from lower to upper quartile values of the data, with a vertical line indicating the population median, whiskers indicate first to 99^th^ percentile. A dashed line is drawn at 0. **(F)***Left:* Summary across all sessions and animals shown in (E). Grid: 6 animals, Mann-Whitney U=320, n_Grid_=n_OV_=18, p=3.82e-07***, two-sided; OV: 6 animals, Mann-Whitney U=3, n_Grid_=n_OV_=16, p=2.70e-06***, two-sided. *Right*: Fraction of cell types (OV or Grid) and discrimination indices in comparisons to a shuffled distribution, which we constructed from assigning Grid or OV labels randomly to cells in our data, assuming that the population ratio is balanced, and the distribution is homogeneous (50% grid and 50% OV cells). We picked 129 cells for each iteration of the shuffling (the average cell number per session for data in (E) and (F), 10,000 shuffling iterations in total). We assigned those shuffled sessions to either (shuffled) Grid or OV animals depending on whether more grid or more OV cells were present in each set. Stars (‘*’) indicate results of the two-sided Wilcoxon signed-rank tests of the data versus the population mean of the shuffled data (* p<0.05, ** p<0.01, *** p<0.001). Grid animals: fraction grid, Z=28, n_Grid_=18, p=0.010*; fraction OV Z=1, n_OV_=18, p=1.53e-05***; discrimination index Z=0, n_Disc.Index_=18, p=7.63e-06***; OV animals: fraction grid, Z=3, n_Grid_=16, p=0.00015***; fraction OV Z=23, n_OV_=16, p=0.018*; discrimination index Z=0, n_Disc.Index_=16, p=3.05e-05***. Boxes in box plots extend from lower to upper quartile values of the data, with a vertical line indicating the population median, whiskers indicate first to 99^th^ percentile.

We subsequently identified 12 out of 15 animals for which object sessions had been run and where on average 10% cells or more crossed grid or OV cell selection criteria (Figure S2H). Depending on whether the fraction of grid or OV cells was higher, these animals were classified as either “Grid” (Figure 3E top row, 3 example animals) or “OV” animals (Figure 3E bottom row, 3 example animals). In these two groups of animals (“Grid” and “OV”), grid and OV cells maintained stable anticorrelated relationships on the population level across multiple sessions (Figure 3E, fraction of OV and Grid cells in relation to all cells, 6 additional animals are shown Figure S3C). In Grid animals, the mean percentage of grid cells (± SD) was 21.2% ± 13.0, whereas OV cells accounted for only 1.2% ± 2.3; in OV animals, the percentage of grid cells was 3.8% ± 2.2, as compared to 17.7% ± 7.8 for OV cells (mean ± SD) (Figure 3F left). To determine the size of the bias in absolute numbers of grid and OV cells, we calculated a discrimination index for each recording (“Disc. Index”, (n_Grid_-n_OV_)/(n_Grid_+n_OV_), Figure 3E and F), ranging from -1 (no grid cells, only OV cells) to 1 (only grid cells). To assess whether the differences in cell fractions and discrimination indices were bigger than expected by chance, we created a shuffled distribution, by labeling a subset of all cells as either grid or OV cells, matching the observed combined frequency of grid and OV cells in our data (932 out of 4399 cells across 34 sessions). We chose a ratio of 50% grid and 50% OV cells, mimicking a balanced population ratio, and drew random subsets from the total cell population that matched the average population size of our recorded sessions (n=129 cells per sample). We then classified these shuffled sets as either “OV” or “Grid”, depending on which cell type was represented more often, as we did for the recorded data. We observed that the differences in percentages of grid and OV cells in the data were more striking than in the shuffled set, both for shuffled “Grid” (Figure 3F right top, mean ± SD, shuffled grid 11.1% ± 2.1, shuffled OV 7.8% ± 1.9, n=4591 shuffles) as well as shuffled “OV” (Figure 3F right bottom, shuffled grid 7.83% ± 1.9, shuffled OV 11.1% ± 2.2, n=4558 shuffles). A similar effect was present in the discrimination indices. Compared to the shuffled distribution, which showed indices close to 0 for shuffled grid and OV sets (Figure 3F right, mean ± SD, grid 0.175 ± 0.118, OV -0.177 ± 0.119), the indices in our data were shifted strongly towards 1 in Grid animals and -1 in OV animals (Figure 3F right, mean ± SD, grid 0.879 ± 0.217; OV -0.617 ± 0.187), indicating substantially stronger biases than expected by chance.

The observed stability of the bias towards either cell type is striking considering that we sampled from varying depths and thus slightly different populations of cells in MEC layer II across days. Most of our recordings were located superficially to layer III, within which the majority of tdTomato expressing cells were found (Figure S1G and Figure S3D for example histology and red cell quantification in one “Grid” (top) and one “OV” (bottom) animal). Due to the large area covered by our implants and hence the difficulty to precisely align postmortem histology and *in vivo* imaging results, we were not able to further pinpoint the exact anatomical location of our imaging FOVs within or around MEC. We noticed however that “Grid animals” were on average implanted close to the parasubicular (PAS) border, while OV animals were implanted either more medially or more laterally than Grid animals (all implant positions in Figure S2H, and analysis below leading up to Figure 6).

In summary we observed that grid cells show a low probability of co-occurrence with OV cells. This trend was maintained across recording days and animals.

### Anatomical separation of grid cells from other functional cell types in MEC

We observed that the fractions of grid and other cell types, particularly OV cells, frequently appeared anticorrelated on the session level (Figure 3). We therefore wondered whether cells that belong to different classes, and which were co-recorded in the same FOV, are randomly spaced relative to each other (intermingle) or occupy separate territories within MEC. The anatomical arrangement of the various cell types in MEC often appeared dispersed across single FOVs (see example cell maps in Figure 4A).

**Figure 4.**
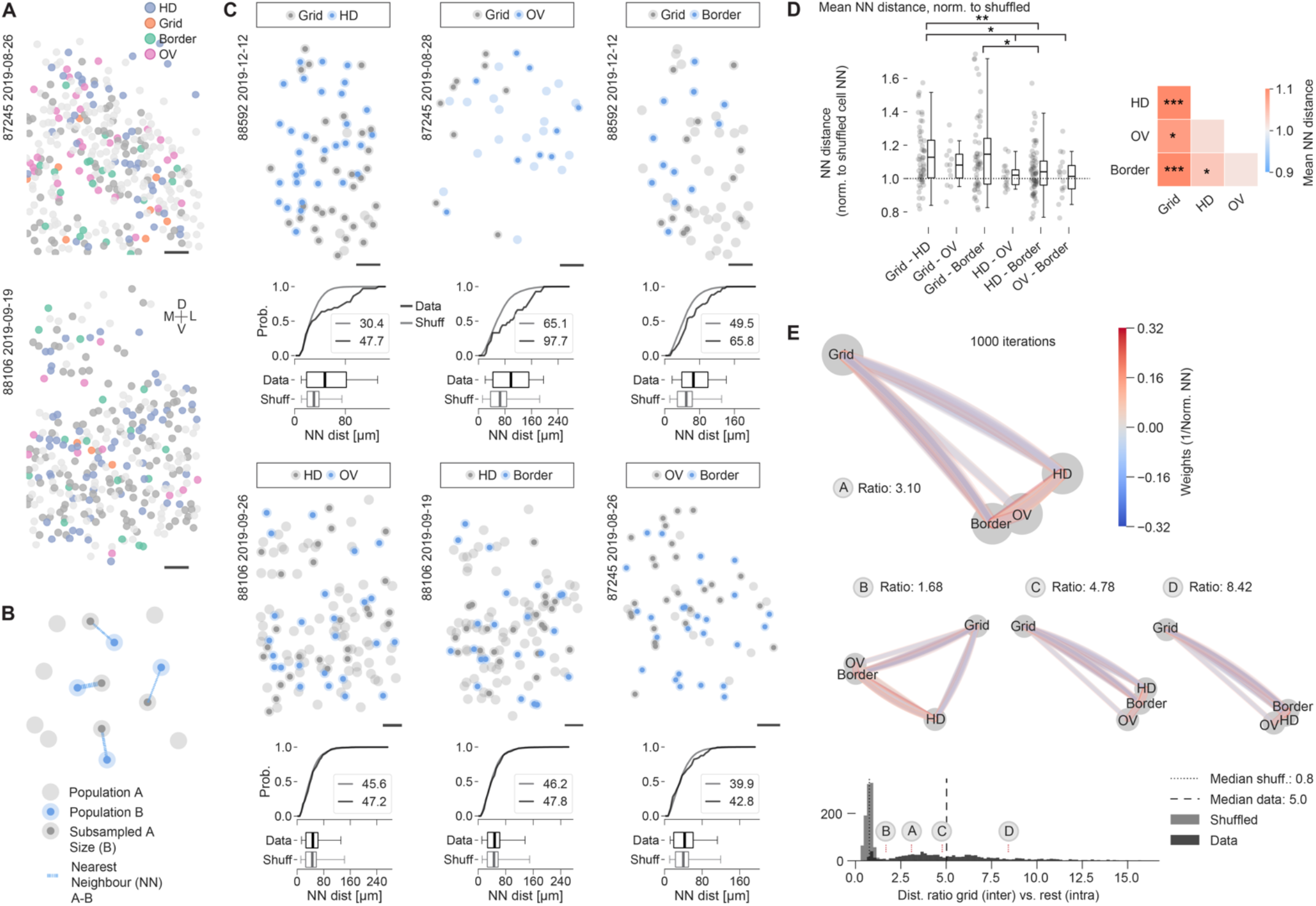
Anatomical separation of grid cells from other functional cell types in MEC. **(A)** Example cell maps of two recordings (taken from OV animals 87245 (322 cells) and 88106 (309 cells)), showing the distribution of HD (blue), Grid (orange), Border (green) and OV (pink) cells. Cells are color labeled if they crossed shuffling cutoff criteria for not more than one cell class. All other co-recorded cells are shown in light grey if they crossed no or dark grey if they crossed more than one cutoff criterion. Tissue orientation is indicated by cross (D-dorsal, L-lateral, V-ventral and M-medial), Scale bar 50 μm. See also Figure S4A for cell maps in pairwise comparisons between cell types. **(B)**Schematic of the nearest neighbor (NN) analyses shown in this figure. The NN distances were extracted from size-matched populations of starter cells (Subsampled A (dark grey) and B (blue) in schematic). Lines indicate NN (A to B) distances (line thickness indicates distance, thinner = longer). **(C)** Representative cell maps and NN distance analyses (underneath cell maps) for six sample pairwise comparisons (Grid/HD, Grid/OV, Grid/Border, HD/OV, HD/Border, OV/Border) across multiple animals (animal name and recording date indicated in top left). Graphs underneath cell maps show, for each example comparison, the cumulative distribution and box plots over nearest neighbor (NN) distances (black) in comparison to a shuffled distribution (shuffled (grey): same population size, but random cell ID). Numbers in insets indicate population averages. Only populations that had a minimum number of five starter cells each were considered. Cells were filtered by anatomical region (MEC) and NN distances were filtered by a minimum distance of 10 μm (cell center distance). For each pairwise comparison only cells crossing either one or the other cell selection threshold were considered. Boxes extend from lower to upper quartile values of the data, with a vertical line indicating the population average, whiskers indicate first to 99^th^ percentile. Scale bar 50 μm. Note the right-shift of data curves compared to shuffled data in comparisons of grid cells and other cell types, i.e., NN distances were larger than expected by chance. There was no corresponding right-shift in comparisons of HD and OV, HD and border cells, and OV and border cells suggesting that these co-localize. **(D)** Normalized mean NN distances over all sessions and animals. NN distances were normalized to a size-matched, shuffled distribution, which was drawn from random cells for each session. Filtered by region (MEC) and minimum cutoff of five cells per cell class. *Left*: Scatter plot showing one dot per dataset. Kruskal-Wallis H-test across all groups: H=14.504, n=6 groups, p=0.013*, Statistics on top of figure indicate results of Mann Whitney U test (summaries shown over results of individual, pairwise comparisons, 95^th^ percentile shuffling cutoffs were used throughout): Grid/HD vs. Grid/OV, U=372, n_Grid/HD_=56, n_Grid/OV_=12, p=0.57 ns; Grid/HD vs. Grid/Border, U=1457, n_Grid/HD_=56, n_Grid/Border_=51, p=0.86 ns; Grid/HD vs. HD/OV, U=627, n_Grid/HD_=56, n_HD/OV_=16, p=0.016*; Grid/HD vs. HD/Border, U=2472, n_Grid/HD_=56, n_HD/Border_=68, p=0.0044**; Grid/HD vs. OV/Border, U=612, n_Grid/HD_=56, n_OV/Border_=16, p=0.027*; Grid/OV vs. Grid/Border, U=282, n_Grid/OV_=12, n_Grid/Border_=51, p=0.68 ns; Grid/OV vs. HD/OV, U=131, n_Grid/OV_=12, n_HD/OV_=16, p=0.11 ns; Grid/OV vs. HD/Border, U=497, n_Grid/OV_=12, n_HD/Border_=68, p=0.23 ns; Grid/OV vs. OV/Border, U=128, n_Grid/OV_=12, n_OV/Border_=16, p=0.14 ns; Grid/Border vs. HD/OV, U=533, n_Grid/Border_=51, n_HD/OV_=16, p=0.067 ns; Grid/Border vs. HD/Border, U=2151, n_Grid/Border_=51, n_HD/Border_=68, p=0.025*; Grid/Border vs. OV/Border, U=539, n_Grid/Border_=51, n_OV/Border_=16, p=0.055 ns; HD/OV vs. HD/Border, U=522, n_HD/OV_=16, n_HD/Border_=68, p=0.81 ns; HD/OV vs. OV/Border, U=135, n_HD/OV_=n_OV/Border_=16, p=0.81 ns; HD/Border vs. OV/Border, U=588, n_HD/Border_=68, n_OV/Border_=16, p=0.62 ns; *Right*: Color coded average of results on the left. Stars indicate results of two-sided one sample t-test against population mean of 1 (p<0.05 *, p<0.01 **, p<0.001 ***); Grid/HD: 12 animals, t=5.364, n_Grid/HD_=56, p=1.67e-06***; Grid/OV: 5 animals, t=2.211, n_Grid/OV_=12, p=0.049*; Grid/Border: 12 animals, t=4.324, n_Grid/Border_=51, p=7.31e-05***; HD/Border: 13 animals, t=2.260, n_HD/Border_=68, p=0.027*; HD/OV: 6 animals, t=1.065, n_HD/OV_=16, p=0.30 ns; OV/Border: 7 animals, t=0.395, n_OV/Border_=16, p=0.7 ns; ns not significant (p>0.05)). See also Figure S4B for quantification of inter vs. intra distance ratios. **(E)** Spring loaded network model with average NN distances acting as weights in between nodes (graph representation, each node represents one cell class). Edges in the graph are color coded by weight (1 / normalized NN distance). The graph at the top shows an example result of one such simulation after 1000 iterations (simulation steps) and more examples with increasing distance ratios (B, 1.68; C, 4.78; D, 8.42; 1000 simulation steps each) across the possible spectrum of results are shown in the middle (see histogram at bottom). The distance ratios depict the inter versus intra ratio of distances (grid cells vs. rest) after running the simulation. Bottom: Results of distance ratios after shuffling weights (n=1000 permutations) versus data (Shuffled: lighter grey, median ratio 0.8 (small-dashed line in histogram), data: darker grey, median ratio 5.0 (large-dashed line in histogram)). See also Figure S4D for examples of shuffled simulation outcomes. Ratio results of the example graphs shown on top are indicated with stippled red lines in the histogram. Note that the first example graph at the top has a ratio that is at the lower end of the distribution, indicating that, on average, more extreme results than the one shown can appear.

To investigate, we quantified the mutual anatomical distances between cell somata over all combinations of cell types at the single session (=single FOV) level (i.e., pairwise comparisons between HD, Grid, Border and OV cells, see example in Figure S4A) after masking out regions that were visibly not part of MEC based on tdTomato expression patterns (see *STAR* methods). We defined starter cell populations (functional cell types above cutoff) for each cell class in each session and quantified nearest neighbor (NN) distances between size matched groups (containing the same number of neurons) from each class (Figure 4B, two starter populations “A” and “B”, minimum no. of cells in each cell class: 5).

We observed that within MEC the somata of grid and HD, grid and OV, and grid and border cells often segregated anatomically from each other (Figure 4C cell maps top row), while HD and OV, HD and Border and OV and Border cells seemed to intermingle (Figure 4C cell maps bottom row). To quantify, we compared the distribution of inter-class nearest neighbor (NN) distances to a size-matched reference distribution, which we constructed by randomly permuting cell IDs for all cells in each session (“Shuffled”). Grid/HD, Grid/OV and Grid/Border distributions frequently appeared shifted towards larger NN distances compared to the reference distribution (cumulative distributions and box plots in Figure 4C) as would be the case if cell classes were anatomically more separated from each other than expected by chance. To compare across all recordings and animals we normalized the average distances to the reference distribution in each recording (Figure 4D). Values above 1 indicate that the average inter-class NN distances (e.g., Grid versus HD NN distances) are larger than those in the reference distribution. We found that the observed trends (examples shown in Figure 4C) were maintained when compared across all recordings, in that Grid/HD, Grid/OV and Grid/Border distributions were shifted above 1 (mean±SD, Grid/HD: 1.13±0.18, Grid/OV: 1.08±0.12, Grid/Border: 1.15±0.24), indicating anatomical separation, while HD/OV, HD/Border, and OV/Border distributions were comparably closer to 1 (mean±SD, HD/OV: 1.02±0.07, HD/Border: 1.04±0.15, OV/Border: 1.01±0.13), indicating anatomical intermingling of cell somata. Grid/HD distributions were statistically different from HD/OV, HD/Border and OV/Border distributions, and Grid/Border distances were statistically different from those of HD/Border (Figure 4D left). Grid/Border, Grid/OV, Grid/HD and HD/Border distributions were significantly different from a population mean of 1 (Figure 4D right). We obtained similar results when analyzing the *inter* (between cell classes) to *intra* (within cell classes) nearest-neighbor distance ratios (Figure S4B) and filtering cells according to different spatial-directional score cutoffs (Figure S4C).

To summarize the results of the multiple pairwise inter-cell class comparisons we constructed undirected graph networks from the data, which allowed us to visualize the pairwise distance relationships in a layout of 2D nodes (Figure 4E, same data as presented in Figure 4D). Each node of these graph networks constitutes one cell class and the weight between each node is determined by the inverse of the normalized distance, such that cell class distances with larger normalized distances have lower weight and those with closer distances higher weight. After random initialization of node positions, we then ran a simulation (“spring loaded model”), which allowed the graph visualization to evolve at each iteration (1000 iterations total) and either draw nodes closer together or push them further apart depending on their weight relationships. Figure 4E (top) shows an example outcome of such a simulation, which shows grid cells separating from all other cell classes. Further examples are shown in the middle row in Figure 4E. We defined a measure of separation as the ratio of the Euclidean distance between the grid node and center of the nodes of all other cell types (*inter* distance) over the mean distance between nodes of all other (non-grid) cell types (*intra* distance) (“Ratio”). Over 1000 shuffled results for which we randomly initialized weights before starting the simulation, the distance ratio distribution had a median close to 1 (0.8, Figure 4E bottom row, more examples in Figure S4D), while simulations run on the real data on average ended up with larger values (median = 5.0, Figure 4E bottom). Increasing the number of iterations yielded even more extreme results (Figure S4E).

Taken together, we provide evidence for anatomical and functional separation of grid cells from all other, known spatially modulated cell classes in superficial MEC.

### Grid cells cluster anatomically

We next wondered whether grid, border, HD and OV cells each exhibit clustering over anatomical space like has previously been reported for grid cells in MEC (Gu et al., 2018; Heys et al., 2014). To address this question, we compared NN distance statistics for groups of starter cells (functional cell types above cutoff, minimum number of starter cells: 15) and reference cells (cells that were not part of the starter cell population, “Ref”) to statistics derived from cells that were picked randomly from all cells in each recording (“All”) (Figure 5A, example cell maps in Figure 5B). We first masked out regions that were visibly not part of MEC (same procedure as for Figure 4). Then, for each recording, we calculated the average nearest neighbor distances over the 5 closest neighbors (NN group 5) of cells in all groups, which were size matched to the starter cell population and normalized to “All” (Normalized NN distances in Figure 5C to E). Values below 1 indicate distances that are smaller than those derived from randomly picked cells (“All”).

**Figure 5.**
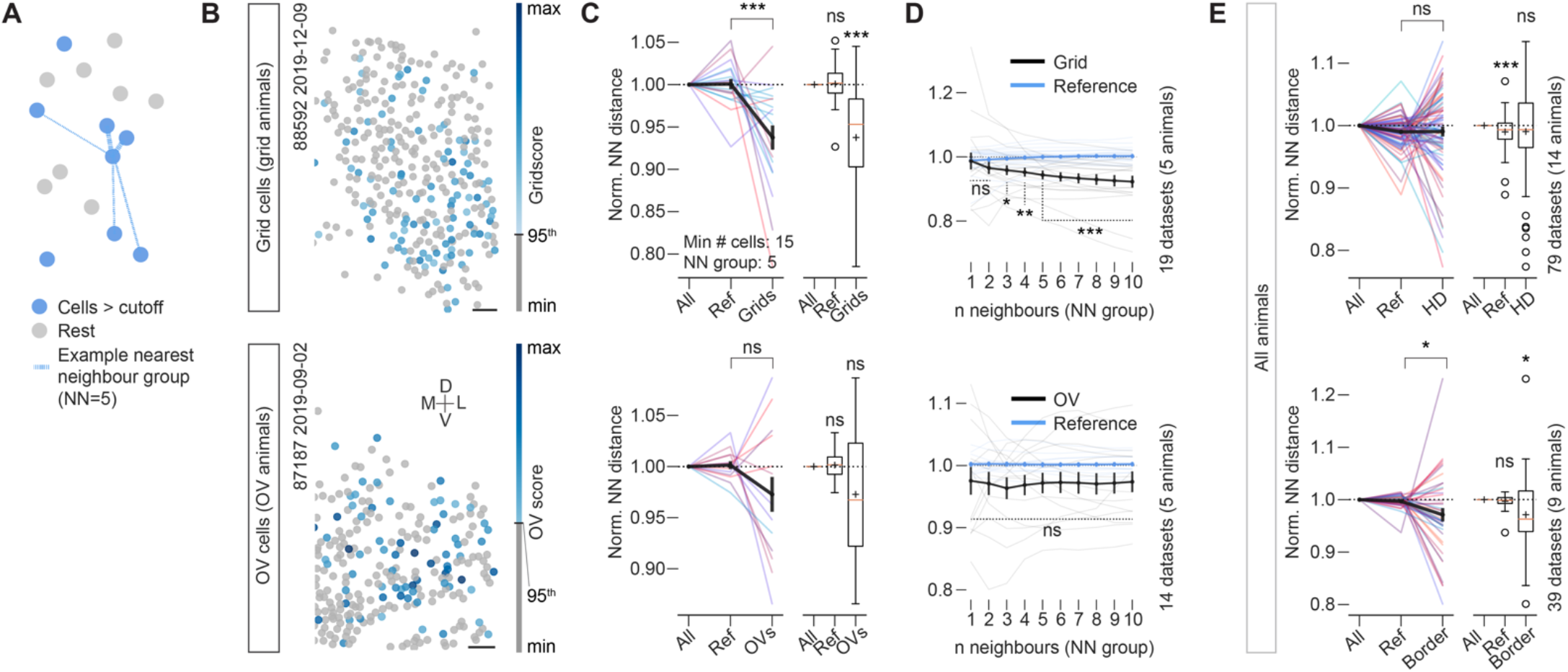
Grid cells cluster anatomically. (B), (C) and (D) show “Grid animals” on top and “OV animals” on bottom. The minimum number of cells above cutoff for each panel was set at 15. Cells were filtered by anatomical region (based on tdTomato signal). **(A)** Schematic explaining the nearest neighbor (NN) analyses shown in this figure. The mean NN distances in (C) and (E) were extracted from groups of 5 starter cells (blue in schematic). Lines indicate distances to 5 nearest neighbors of example cell in center. This group size (“NN group”) is systematically varied in (D). **(B)** Two sessions in two example animals in which multiple grid cells (“Grid animal”, top) or multiple OV cells (“OV animal”, bottom) were recorded (color coded by gridscore or OV score). See also Figure S5B. Cells that did not meet cutoff criteria (grid: 95^th^ percentile shuffling) are color coded in grey, animal number and recording date are indicated in top left, anatomical orientation indicated in bottom left (M-medial, L-lateral, V-ventral, D-dorsal). Scale bars 50 μm. **(C)** Mean NN distances are normalized to a distribution of cells randomly picked out of all cell locations available for each session and of the same population size as the population of functional cell types under investigation (“All”). A second comparison is shown (reference “Ref”), which consists of cells that were explicitly not part of the cell population under investigation (i.e., non-grid or non-OV cells; same population size as grid / OV cells). Left: Each line indicates one recording session and colors represent animals. Right: Boxplots: boxes extend from lower to upper quartile values of the data, with an orange line at the median, whiskers indicate 1^st^ to 99^th^ percentile, outliers are shown as open black circles, black plus signs indicate mean. Grid: data over 5 animals, Mann-Whitney U=308, n_Ref_=n_Grid_=19, p=0.00021*** two-sided; Wilcoxon signed-rank test (against 1.): Ref: Z=90, n_Ref_=19, p=0.86 ns, Grid: Z=9, n_Grid_=19, p=0.00013***; OV: data over 5 animals, Mann-Whitney U=131, n_Ref_=n_OV_=14, p=0.14 ns two-sided; Wilcoxon signed-rank test Z=30, n_OV_=14, p=0.17 ns two-sided. Thick black lines indicate Mean ± SEM; ns: not significant (p>0.05). See also Figure S5A for results across various shuffling cutoffs and Figure S5C to F for analyses of all pairwise distances instead of NN groups. **(D)** Mean nearest neighbor (NN) distance over varying numbers of neighbors (“NN group”, a number of 5 was used in (C) and (E)) for grid cells in “grid animals” (top) and OV cells in “OV animals” (bottom). “Reference” is picked as described for (C) (“Ref”). Thin black and blue lines show single recordings and thick lines show group average and SEM. Data was normalized to a subsampled population of all cells in each recording (as in (C)). The clustering in grid cells is not an effect of “doublets” / “triplets” / etc. but evolving and stable over large numbers of cells under consideration. Significance (p<0.05*, p<0.01**, p<0.001***) indicates the result of the two-sided Mann-Whitney U test Data vs. Reference. Vertical labeling towards the right shows the total number of animals and sessions that were used in each comparison (applies to both (C) and (D)). Mann-Whitney U-test results: Grid vs. Ref, n_Ref_=n_Grid_=19; NN=1 U=147, p=0.34 ns; NN=2 U=115, p=0.058 ns; NN=3 U=103, p=0.025*; NN=4 U=89, p=0.0079**; NN=5 U=66, p=0.00087***; NN=6 U=53, p=0.00021***; NN=7 U=49, p=0.00013***; NN=8 U=50, p=0.00015***; NN=9 U=48, p=0.00012***; NN=10 U=46, p=9.15e-05***; OV vs. Ref: n_Ref_=n_OV_=14; NN=1 U=77, p=0.35 ns; NN=2 U=66, p=0.15 ns; NN=3 U=65, p=0.14 ns; NN=4 U=69, p=0.19 ns; NN=5 U=72, p=0.24 ns; NN=6 U=65, p=0.14 ns; NN=7 U=67, p=0.16 ns; NN=8 U=64, p=0.12 ns; NN=9 U=66, p=0.15 ns; NN=10 U=68, p=0.18 ns; ns: not significant (p>0.05). **(E)** Normalized, mean nearest neighbor (NN) distances as in (C) for head direction (HD) cells (top, 95^th^ percentile shuffling cutoff) and border cells (bottom, 95^th^ percentile shuffling cutoff) including all animals. HD: data over 14 animals, Mann-Whitney U=2887, n_Ref_=n_HD_=79, p=0.42 ns two-sided; Wilcoxon signed-rank test (against 1.): Ref: Z=894, n_Ref_=79, p=0.0008*** two-sided, HD: Z=1467, n_HD_=79, p=0.58 ns, two-sided. Border: data over 9 animals, Mann-Whitney U=1005, n_Ref_=n_Border_=39, p=0.015* two-sided; Wilcoxon signed-rank test (against 1.), Ref: Z=305, n_Ref_=39, p=0.24 ns two-sided, Border: Z=222, n_Border_=39, p=0.019*, two-sided; ns: not significant (p>0.05).

Similar to previous reports (Gu et al., 2018; Heys et al., 2014) we observed that the NN statistics of grid cells indicated strong clustering in “grid animals” (Figure 5C top row, “Grids” mean ± SEM: 0.94 ± 0.01, same group of animals as in Figure 3), which was maintained across cell inclusion thresholds (95^th^ percentile shuffling cutoff in Figure 5; 99^th^ percentile shuffling cutoff in Figure S5A left: “Grids” mean ± SEM: 0.92 ± 0.02). When we varied the NN group sizes from 1 to 10 cells (Figure 5D top), clustering became statistically significant when groups were expanded beyond two neighbors and stayed robust over successively larger NN group sizes (Figure 5D). This indicates that the observed clustering was not a consequence of doublets or triplets of cells, but stable over large groups of neurons.

A similar trend was observed for OV cells in “OV animals” (same group of animals as in Figure 3), which did however not reach statistical significance and was unstable over varying group sizes (Figure 5C and D bottom row, “OVs” mean ± SEM: 0.97 ± 0.02). Border and HD cells exhibited distance relationships that were less conclusive overall. While HD cells showed an overall dispersed arrangement (Figure 5E top, statistics over all animals; “HD” mean ± SEM: 0.99 ± 0.01, Figure S5A middle (99^th^ percentile cutoff): 1.00 ± 0.01), border cells did show some degree of clustering (Figure 5E bottom, statistics over all animals; “Border” mean ± SEM: 0.97 ± 0.01), but it appeared less stable over varying cutoffs (Figure S5A right (99^th^ percentile cutoff): 0.98 ± 0.02). Similar results were obtained when not nearest neighbor distances over groups of cells with fixed number of neighbors but all pairwise distances between starter cells were considered (median pairwise distances over all cell pairs, Figure S5B to F). For this analysis we also varied the cutoff for the minimum number of starter cells (Figure S5D) and subsequently included all animals in the analysis (Figure S5E, no pre-selection for “grid” or “OV” animals), neither of which changed the results.

All in all, we were able to confirm previous reports of clustering in grid cells and extended this analysis over all other spatially modulated cell types. While some cell classes like OV and border cells did show trends for decreased nearest neighbor distances compared to a reference population, none of the other cell classes exhibited anatomical clustering as strongly as grid cells.

### Grid, HD, and border cells cover stable anatomical territories along the MEC / PAS border

Upon closer investigation of our implantation sites and tdTomato labeling patterns *in vivo* we noticed that animals in which we recorded an abundance of grid cells were often implanted quite medially, that is, close to the boundary between MEC and the more medially located parasubiculum (PAS). In these animals (“MEC/PAS animals”), which we frequently recorded over weeks, clear and stable demarcations in the tdTomato channel were discernible, with cells in superficial MEC showing strong labeling in this channel compared to neighboring, more medial structures (example in Figure S6A, overview of all implant positions and tdTomato border visibility in Figure S2H right). This matches descriptions of anatomical projection patterns from superficial MEC to ipsilateral hippocampus (Witter et al., 2017). Moreover, we noticed that the fractions of functional cell types stayed relatively stable across recording days, with grid, HD and border cells showing roughly similar population ratios (Figure S6B), indicating that our recording locations and the recorded networks themselves remained stable.

We thus wondered if and how the observed anatomical patterns overlapped with the distribution of functional cell types in these regions. To investigate this question, we first aligned the FOVs of as many recordings as possible for each animal to each other based on anatomical landmarks (mean number FOVs per animal: 8.6, n=15 animals) (Figure 6A). This way we were able to create composites with high anatomical congruence (Figure 6A middle, structural similarity index measure, SSIM), aligned across multiple recordings that were spread over many days in each animal (see example in Figure 6A right). We manually annotated MEC and neighboring structures in every FOV (overview in Figure S6C left) and created topographic tuning maps from all neurons by first binning each cell’s tuning strength (e.g., its grid score, etc.) in anatomical space, and then projecting those binned maps over all aligned sessions (difference between first and last session, mean ± SD: 22.1 ± 12.9 days, n=15 animals). Examples can be seen in Figure 6B for one MEC/PAS animal (10 aligned sessions, boundary is indicated by a stippled gray line in each topographic tuning map). In animals in which this clean anatomical boundary was discernible, grid cells cluster on the MEC side (more examples shown in Figure S6C). Sharp transitions occurred towards neighboring PAS in which border and HD cells dominated (Figure 6B). To assess whether these functional transitions were more abrupt than expected by chance and hence whether stable anatomical territories were formed by either cell type, we calculated a global autocorrelation statistic (Moran’s I, Figure 6B and Figure S6C), which we compared to shuffled distributions of the same data. The shuffled distributions were constructed from projections of the same underlying cell position data after scrambling the tuning and ROI associations. These shuffled topographic tuning maps thereby mimicked a “salt and pepper” organization across the tissue. The analysis showed that the anatomical expression patterns of functional cell types across MEC and PAS were more structured than expected by chance (Figure 6B bottom row, see Figure S6C for 5 animals that were implanted at a similar location). In addition, grid modules appeared clustered according to gradients described in the literature (Stensola et al., 2012), with smaller spaced grid cells located more dorsally than larger spaced ones (see Figure S6D to F for an example and procedure of grid module extraction).

**Figure 6.**
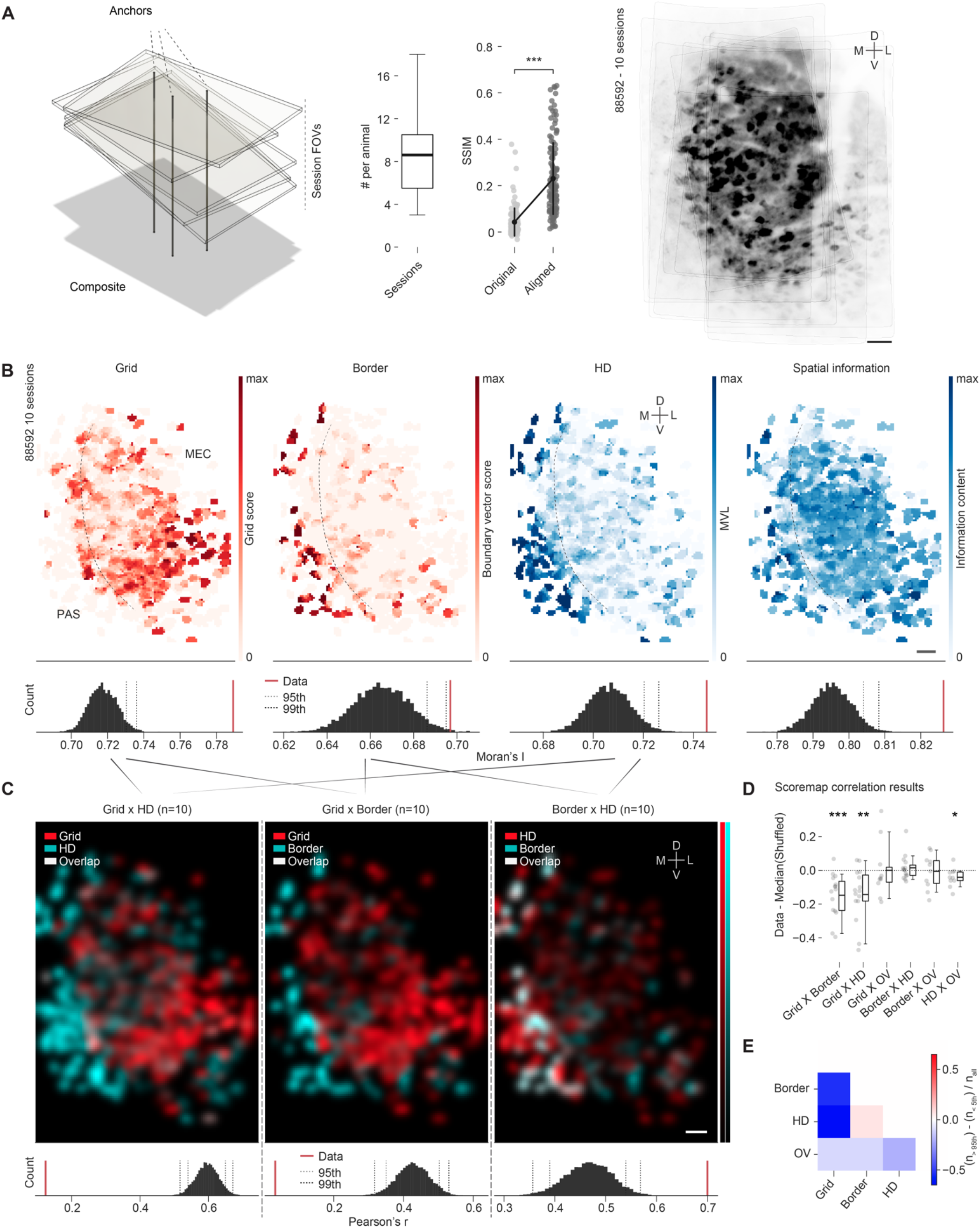
Functional cell types cover distinct and stable anatomical territories. **(A)** Left: Schematic of alignment process for multi-session composites. Multiple fields of view (FOVs) from one animal can be stitched into a composite after defining anatomical anchors (salient landmarks like cells, blood vessels, etc., which are present in multiple recordings) and subsequent affine transformation over those anchors. Middle left: Number of sessions per animal, which were aligned (mean=8.6, n=15 animals). Middle right: Structural similarity index measure (SSIM) comparison between overlapping regions in raw (“Original”) and aligned image pairs (“Aligned”) shows that alignment was successful across FOVs (Mann-Whitney U=1373, n_Original_=n_Aligned_=134, p=4.20e-33***). Right: Composite (average projection) of maximum intensity images for 10 sessions from one animal after alignment. Scale bar 50 μm. **(B)** Topographic tuning maps were created by binning ROI coordinates after alignment (see (A)) and creating average projections across all recordings for specific tuning properties (e.g., grid score). Four such example tuning maps, color coded by score, are shown for one animal (same as in (A) right) for (left to right): grid score (“Grid”), boundary vector score (“Border”), head direction tuning expressed as mean vector length (MVL, “HD”) and spatial information content (“Spatial information”). A sharp border in the composition of functional cell types (dashed line) is visible at the border between medial entorhinal cortex (MEC, towards right) and parasubiculum (PAS, towards left) (derived from tdTomato signal). Bottom: Distribution of shuffled Moran’s I values for each tuning map and actual values (“Data”, red). 95^th^ and 99^th^ percentile cutoffs are shown as dashed lines. All topographic tuning maps exhibit more clustering (i.e., internal structure) than expected by chance (> 99^th^ cutoff). Scale bar 50 μm. **(C)** Overlay and correlation of topographic tuning maps for different tuning properties shows anatomical exclusion of the territories of grid vs. HD and border cells. Maps were first smoothed with a gaussian kernel with sigma 2 bins. The number of combined sessions in each example is shown on top (n). Diagonal lines indicate which maps were combined and smoothed (composites in (B)). Bottom: Pearson’s correlation (Pearson’s r) between the different tuning maps, compared to a shuffled distribution. To create the shuffled distribution, randomized tuning maps were compared to each other, which were themselves created by picking a shuffled cell map for each session (same cell positions, shuffled cell ID) and creating a projection as for the actual data. Actual values are shown in red (“Data”), and 95^th^ and 99^th^ percentile cutoffs are shown as dashed lines. Scale bar 50 μm. **(D)** Topographic tuning map correlation results over all animals (dots indicate animals). Data are shown as the difference of the actual Pearson’s r value (“Data”) and the median of the shuffled distribution of Pearson’s r values (“Shuffled”). Kruskal Wallis H-test over n=6 groups p=0.0012**, H=20.18. Statistics on top of figure indicate statistically significant (p<0.05) results of two-sided Wilcoxon signed-rank test against mean of zero: GridxBorder: Z=4, n_GridxBorder_=15, p=0.00043***; GridxHD: Z=9., n_GridxHD_=15, p=0.002**; GridxOV: Z=21, n_GridxOV_=10, p=0.56 ns; BorderxHD: Z=55, n_BorderxHD_=15, p=0.80 ns; BorderxOV: Z=25, n_BorderxOV_=10, p=0.85 ns; HDxOV: Z=8, n_HDxOV_=10, p=0.049*. **(E)** Summary of topographic tuning map correlation results. Data are shown as a color-coded representation of the difference in the number of results that are above the 95^th^ percentile shuffling cutoff, and the number of results that are below the 5th percentile cutoff over all sessions (same data as analyzed in (D). GridxBorder: -0.53; GridxHD: -0.67; GridxOV: -0.1; BorderxHD: 0.07; BorderxOV: -0.1; HDxOV: -0.2). Grid / Border as well as Grid / HD territories are most strikingly anti-correlated compared to the rest (blue coloring).

Furthermore, the territories of grid, border, and HD cells appeared spatially anticorrelated at the MEC/PAS boundary (Figure 6C, bottom row shows Pearson’s correlation between smoothed topographic tuning maps compared to shuffled distribution). We compared the strength of this effect across all animals, including those that were not implanted close to the PAS boundary, by subtracting the median of all shuffled Pearson’s correlation values from the data (Figure 6D, values below zero indicate that expression patterns were more anticorrelated than expected by chance). To summarize these data, we compared the number of results above the 95^th^ percentile shuffling distribution cutoff, i.e., topographic tuning maps that were most strongly correlated, to those that were below the 5^th^ percentile shuffling distribution cutoff, i.e., those that were most strongly anticorrelated across all cell classes (discrimination ratio, Figure 6E). In line with our observations of functional territories in MEC/PAS animals and our analyses of cell soma distribution patterns on the single session level (Figure 4), grid cell territories were most stably distinct from HD and border cell territories, while OV, border and HD territories appeared to overlap more (Figure 6D, mean ± SD: Grid×Border -0.15±0.13 (15 animals), Grid×HD -0.14±0.15 (15 animals), Grid×OV 0.00±0.16 (10 animals), Border×HD 0.01±0.07 (15 animals), Border×OV -0.01±0.10 (10 animals), HD×OV -0.04±0.05 (10 animals)). Because of the low co-expression level of grid and OV cells in the FOVs that were accessible to us in this study, the analysis of anatomical territories was less powerful in the Grid×OV case (Figure 6D and E), although our previous analyses predict strong anticorrelation of grid and OV distribution patterns at larger anatomical scales. Future studies that make use of further improvements that are described in the accompanying Resource article (Zong et al.), enabling both the enlargement of single session FOVs as well as the systematic shifting of FOVs between sessions, may extend the present investigations to even larger anatomical spaces.

The robustness of the observed effects indicates that functional cell types occupy anatomically stable territories over time, with the allocation of grid and HD/border networks on either side of the MEC/PAS boundary leading to the most apparent and stable topographical distribution differences.

## Discussion

We have shown that with a new generation of 2P miniscopes we can, for the first time, systematically investigate the topographical arrangement of all known spatially modulated cell types in the superficial layers of MEC and adjacent PAS. While *in vivo* calcium imaging is ideally suited to address these questions, previous endeavors in this direction have largely been held back by being limited to imaging through either miniature 1P microscopes (1P miniscopes), or benchtop 2P microscopes. In 1P miniscopes the optical sectioning capability is much more limited than with 2P imaging, which renders them susceptible to out of focus fluorescence that can interfere with the identification and analysis of individual cells in densely labeled and active networks like the MEC. However, their light weight and flexible connection cable enable mice to move freely in open environments (Ghosh et al., 2011; Sheintuch et al., 2020; Sun et al., 2015; Ziv et al., 2013). On the other hand, benchtop 2P microscopes allow for high-resolution, high signal-to-noise ratio recordings, but require animals to be head-fixed, which impedes the functional expression of most spatial cell types due to the absence of any vestibular input (Minderer et al., 2016). 2P miniscopes used in our study combine the best of both worlds: Their light weight and flexible connection cable is on par with that of previous 1P miniscopes and their spatial resolution approaches that of 2P benchtop setups. Our data demonstrate that 2P miniscopes enable the simultaneous recording of well over a hundred cleanly isolated cells at high SNRs while mice engage in unrestrained, exploratory behavior in large open field arenas over tens of minutes. Improvements to this technology are ongoing and are reported in detail in the accompanying Resource article by Zong et al. Future studies using these upgraded 2P miniscopes will benefit from further increases in the total number of recordable cells per FOV as well as the ability to systematically expand surveys over multiple layers and multiple neighboring anatomical regions in the same animal.

We report that grid and object vector cells in MEC display low co-occurrence on the anatomical scales that were accessible to us in this study. Animals segregate into groups with either high numbers of grid cells or high numbers of OV cells per FOV. This is unlikely to be a result of sampling different layers within MEC, since our recordings were mostly obtained from layer II or at the border between layer II and III and even if they varied slightly from day to day, the observed anticorrelations on the population level between OV and grid cells remained stable across recordings and over days. Moreover, we did not find any correlation between the fraction of OV or grid cells and numbers of tdTomato positive cells, the latter of which were more numerous in layer III than layer II in our study. Together, this suggests that grid and OV cells occupy largely separate territories in the parahippocampal region and could indicate that the tuning properties of grid and OV cells emerge in distinct anatomical sub-networks. The segregation of OV and grid cells might hereby reflect the different affordances of underlying network computations. These involve, in the case of grid cells, the formation of internally generated spatial firing patterns anchored to the overall layout of the environment the animal is in (Gardner et al., 2021; Hafting et al., 2005), and, in the case of OV cells, the formation of firing fields that anchor to and move flexibly with objects within that same environment (Høydal et al., 2019). Similar dissociations have previously been explored in the context of hippocampal neurons that can appear anchored to either local or distal cues in “double rotation” experiments, which aim to bring the two reference frames into direct conflict with each other (Shapiro et al., 1997; Tanila et al., 1997). Various degrees of anchoring preferences within the same population of recorded hippocampal cells (Knierim, 2002; Neunuebel et al., 2013; Shapiro et al., 1997; Tanila et al., 1997; Zinyuk et al., 2000) might partially be inherited from upstream grid (*distal* cue anchored) or OV (*local* cue anchored) cell networks in the parahippocampal region. Cell populations that lead to the emergence of grid or OV cells and, subsequently, tuning preferences of downstream hippocampal neurons might thus feature differential connectivity motifs. These motifs might bias populations of neurons in MEC and neighboring regions towards stably supporting either grid or OV cells. Connectivity divergences like this have been proposed previously for networks supporting the emergence of aperiodic spatial cells and grid cells in MEC (Miao et al., 2017), which might be relevant if OV cells are at least to some extent contained within the miscellaneous class of aperiodic spatial cells (Andersson et al., 2021). Although we were not able to resolve the exact number and location of grid or OV territories in this study, future studies using technological advances described in the accompanying *Resource* article, featuring 2P miniscopes with even larger FOVs (Zong et al., accompanying paper), will extend the anatomically observable areas and thereby help to elucidate these large-scale organizational principles further.

The somata of grid cells cluster within MEC, which is in line with previous reports (Gu et al., 2018; Heys et al., 2014). This topographical pattern seems to go along with the anatomical separation from various other cell classes, visible within MEC and in between MEC and the neighboring parasubiculum. The dense anatomical packaging of grid cells may lead to elevated pairwise connectivity between them (Udvary et al., 2020), which is to be expected if grid cells are organized in continuous attractor networks (CAN) with dense, recurrent connectivity motifs (Burak and Fiete, 2009; Couey et al., 2013; Fuhs and Touretzky, 2006; McNaughton et al., 2006). Here we extend these measurements to all other spatially modulated cell classes and observe that, while trends for anatomical clustering do exist in some cell classes, namely OV and border cells, only grid cells seem to stably aggregate in groups of three or more cells. While we cannot rule out that for example OV and border cells might display more apparent clustering in different anatomical regions and over larger samples, this result could indicate that HD, border and OV cells form more loosely organized attractors or might instead partially inherit their intrinsic dynamics from upstream networks, which would in turn alleviate the need for close anatomical packing within MEC itself. It may however also indicate that these cell types are embedded within the same densely interconnected network. Indeed, we observe that OV, border and HD cells appear anatomically intermingled, resembling a “salt and pepper” organization. This more homogeneous distribution, and thereby the anatomical closeness across compared to within types, might favor strong communication between cell types (Udvary et al., 2020), which may be more difficult to maintain if cells were arranged in anatomically isolated clusters instead. This might be necessary for the emergence and maintenance of firing patterns in these three cell types. In line with this argument, recent studies (Bicanski and Burgess, 2018; Gofman et al., 2019; Peyrache et al., 2017) report that the combination of allocentric head direction coding and egocentric information in relation to environmental boundaries might have an important role in forming border and boundary vector cell firing fields. Border and HD cells might thus benefit from anatomical closeness and thus potentially dense information exchange between each other. Similar mechanisms might require object vector cells to maintain anatomical proximity to cells tuned to head direction and environmental boundaries. Future studies that combine 2P miniscope imaging with single cell optogenetic stimulation protocols in functionally identified neuronal subtypes might be able to elucidate these connectivity motifs.

Our data indicate that grid neural networks do form largely isolated entities in the superficial layers of MEC, which may isolate them from direct connectivity with other spatially modulated cell classes. In line with this, we found reliable “hotspots” of grid cells in MEC at the border to PAS across animals, which appeared to span large parts of the imaged FOVs. PAS seemed to be populated with mostly HD and border cells and fewer, more scattered grid cells, which is in line with previous reports (Boccara et al., 2010; Tang et al., 2016). Although grid cells in MEC are in close contact with HD and border cells in superficial PAS, they respect a strikingly clean anatomical boundary that is visible across animals and, within animals, remains stable over multiple days and weeks. Together with the observation of anatomical segregation within MEC, this hints at a relatively rigid, developmentally controlled process, which leads to the formation of highly structured intrinsic networks in the parahippocampal region. The anatomical segregation of clustered grid networks on the one hand and intermingled border and head direction networks on the other hand might be seeded during development of the parahippocampal network, in line with observations that the maturation of grid cells is protracted compared to both HD and border cells (Bjerknes et al., 2014, 2015; Langston et al., 2010; Wills et al., 2010). HD, border and potentially OV cells, for which the developmental time course has not yet been explored, might form early, intermingled subnetworks that foster the establishment of grid networks. Future studies employing 2P miniscope imaging in combination with clever labeling of genetically accessible subpopulations might yield insights into how this internal organization is orchestrated during development, specifically how functional arrangements follow known anatomical developmental patterns in this region (Baram et al., 2019; Donato et al., 2017).

## Supporting information

Key Resources Table

## Acknowledgements

We thank M.P. Witter for help with evaluation of recording locations, and Simon Ball, A.M. Amundsgård, K. Haugen, K. Jenssen, E. Kråkvik, Ingvild Ulsaker-Janke, and H. Waade for technical assistance. We thank R. N. Raveendran and the Viral Vector Core Facility of the Kavli Institute for Systems Neuroscience (NTNU) for viral reagents. We also thank Debora Ledergerber for providing the original MATLAB code for the calculation of boundary vector scores in Figure 6, which we adopted and modified in our study. The work was supported by a Synergy Grant to E.I.M. and Yoram Burak from the European Research Council (‘KILONEURONS’, Grant Agreement N° 951319), grants from the Research Council of Norway (FRIPRO to E.I.M., grant number 286225; Centre of Excellence to M.-B.M. and E.I.M., grant number 223262; National Infrastructure to E.I.M. and M.-B.M., NORBRAIN, grant number 295721), a Research Council of Norway (RCN) FRIPRO grant to M.-B.M. (grant number 300394), the Kavli Foundation (M.-B.M. and E.I.M.), and a direct contribution to M.-B.M. and E.I.M. from the Ministry of Education and Research of Norway, support by the Max Planck Society (T.R. and T.B) and the German Research Foundation (DFG/CRC870_A07, Project-Nr. 1188 03580; T.R. and T.B.), a European Research Council Starting Grant ERC-ST2019 850769 (F.D.) and NSFC No. 81925022, 92054301, 31821091 (L.C.), and Beijing Natural Science Foundation Key Research Topics No. Z20J00059 (L.C.).

## Author Contributions

H.A.O., T.B., M.-B.M. and E.I.M. planned the project and designed experiments; H.A.O. and W.Z. built the imaging setup with input from T.R. and T.B.; W.Z. designed and built improved version of miniscope; H.A.O., W.Z. and R.I.J. performed experiments; F.D. helped with surgical methods; L.C. and H.C. provided miniscope technological support; H.A.O. developed and performed the analyses; H.A.O., M.-B.M. and E.I.M. interpreted data; H.A.O. visualized data; H.A.O wrote the paper; H.A.O., E.I.M. and M.-B.M. edited the paper, with input from all authors; E.I.M., M.-B.M., T.B. and H.C. supervised the project; E.I.M., M.-B.M. and T.B. obtained funding.

## *STAR* METHODS

### RESOURCE AVAILABILITY

#### Lead contact

Further information and requests for resources and data should be directed to and will be fulfilled by the lead contact, **Edvard I. Moser**.

#### Materials availability

This study did not generate new unique reagents.

#### Data and code availability

The data for each figure in this manuscript will be available as of the date of publication in a standardized format accessible in Python. DOIs will be listed in the key resources table. Microscopy data reported in this paper will be shared upon request.

All original code used in generation of figures has been deposited on github and will be publicly available as of the date of publication. DOIs will be listed in the key resources table.

Any additional information required to reanalyze the data reported in this paper is available upon request.

## EXPERIMENTAL MODEL DETAILS

### Animals

All experiments were performed in accordance with the Norwegian Animal Welfare Act and the European Convention for the Protection of Vertebrate Animals used for Experimental and Other Scientific Purposes, permit numbers 18013, 6021 and 7163. Only adult male mice (age >3 months) were used. C57BL/6JBomTac mice (“wild-type”, Cat#B6JBOM; RRID: IMSR_TAC:b6jbom, Taconic) were housed in social groups of 2-6 individuals per cage under a 12h light/12h darkness schedule, in a temperature- and humidity-controlled vivarium. Food and water were provided ad libitum. The animals were housed under conditions free from specific pathogens according to the recommendations set by the Federation of European Laboratory Animal Science Associations (FELASA working group on revision of guidelines for health monitoring of rodents and rabbits et al., 2014).

For most *in vivo* imaging experiments, GCaMP transgenic mice were used, which express GCaMP6s ubiquitously in excitatory cells under control of the CaMKII promoter (Wekselblatt et al., 2016) (Camk2a-tTA; tetO-G6s). Due to observed physiological abnormalities that we observed in old (age >11 months) transgenic animals, only results from transgenic animals that were younger than 10.5 months at the time of recording are reported in this study (age at recording mean ± SD 29.57 ± 5.82 weeks, n=103 sessions).

## METHOD DETAILS

### Viruses

In a subset of animals, a virus expressing GCaMP6m (AAV1-syn-GCaMP6m, titer: 3.43e13 GC/ml, Cat#AV-1-PV2823, UPenn Vector Core, University of Pennsylvania, USA) was injected into wild-type animals. Most animals (transgenics and wild-types) were injected with retrograde AAV (Tervo et al., 2016) carrying tdTomato into the hippocampus (AAVretro-CAG-tdTomato, #59462-AAVrg, Addgene).

### Surgeries

For all surgeries anesthesia was induced by placing the subjects in a plexiglass chamber filled with isoflurane vapor (5% isoflurane, IsoFlo, Zoetis, in medical air, flow of 1 liters/minute). Surgery was performed on a heated surgery table (38°C) and air flow was kept at 1 liters/minute with 1.5–3% isoflurane as necessary based on depth of anesthesia, as determined by physiological monitoring of vital signs (breathing and heartbeat). After each procedure, subjects were allowed to recover in a heated chamber (33 °C) until they regained complete mobility and alertness.

### Virus injection into the hippocampus

On the day of surgery, adult mice (P60-P120) were anesthetized, and analgesics were provided (Rymadil, Pfizer, intraperitoneal injection, 5 mg/kg or Metacam, Boehringer Ingelheim, 5 mg/kg; Temgesic, Indivior 0.05-0.1 mg/kg subcutaneous injection, 0.05 mg/kg; Marcain, Aspen, local subcutaneous injection, 1-3 mg/kg). Eyes were protected from drying out by applying eye ointment (Simplex, Actavis). Most animals received an injection of AAVretro-CAG-tdTomato into the hippocampus to retrogradely label projections from ipsilateral, superficial layers of MEC. For this, two drill holes (drill bit: Cat#1RF HP 330 104 001 001 005 from Hagen and Meisinger, Germany) were made over the left hemisphere at [mm, measured from bregma] AP 2.1, ML 1, targeting dentate gyrus (DG), and AP 2.1 mm, ML 2.1, targeting CA3; for animals 97045 and 97046 AP was changed to 1.7. After drilling and exposing the brain surface, the drill holes were immediately covered with drops of saline (NaCl 9 mg/ml, B. Braun Medical). The virus was injected at the same coordinates as the drilled holes, at DV 1.8 and DV 1.7 for targeting the DG and CA3, respectively, at a rate of about 50-70 nl per minute via pulled glass micropipettes (Cat#504949, World precision instruments). The total injection volume ranged from 150 to 300 nl per injection site. After each injection was completed, the glass pipette was left in place for 10 minutes to give the virus time to diffuse before retraction. The drill holes were covered with UV curable cement (Venus Diamond Flow, Kulzer).

Virus injections into the hippocampus were typically combined with MEC virus injections and implantations (see below). In rare cases where these procedures were split over weeks, the skin was sutured (Supramid DS 13, Resorba Medical, Germany), and the animal allowed to recover in a heated chamber (33°C, 30-90 minutes).

### Virus injection into MEC and GRIN / prism implantation

In most cases the injections described above (AAVretro-CAG-tdTomato, hippocampus) were combined with virus injections and chronic implantation of GRIN / prisms into MEC. To gain access to MEC a large circular craniotomy (diameter ∼ 4 mm) was made over the left hemisphere. For this, the skull was carefully leveled using bregma and lambda as measurement points and two reference drill points were created at [mm] 3.3 and 3.8 ML on the lambdoid suture. Then a craniotomy was drilled around the reference points, covering mostly the left interparietal bone, with about one fourth of it stretching over to the left parietal bone. The bone was carefully lifted, exposing the brain surface and transverse sinus. The implantation side was frequently irrigated with saline (NaCl 9 mg/ml, B. Braun Medical) and small bleedings were stopped with highly absorbent sponges (Sugi Eyespear, Cat#30601, Questalpha, Germany). For virus injections into MEC, a Hamilton syringe (World Precision Instruments, microliter syringe #75, Hamilton Company) was used. Two injection sites were chosen at [mm] 3.2-3.3 ML and 3.8 ML with the frontal edge of the transverse sinus acting as reference for these coordinates: The medial injection site was offset about .25 mm (towards anterior) and the lateral about .2 mm and the syringe was angled at 9-10 degrees (tip towards posterior). Small incisions were then made in the dura with a syringe needle just underneath where the syringe tip interfaced with the brain surface and the injection needle was inserted and slowly progressed until it touched dura or bone. The needle was then retracted about .1-.2 mm to target superficial layers of MEC and 400-500 nl virus was injected at a rate of 80 nl/min across two depths, spaced about 150-250 μm apart. After each injection the needle was left in place for 10 minutes before retraction.

The craniotomy was then protected by applying Kwik-Cast (World Precision Instruments) and the rest of the exposed skull was covered with a bonding agent (OptiBond All-In-One, Kerr). After removal of Kwik-Cast, the dura was sliced along the transverse sinus and a custom implant holder, attached to a stereotactic micromanipulator (1760, Kopf, CA, USA), clamping a customized 1 mm diameter GRIN (gradient-index) lens attached to a 1 mm square prism (total length approximately 4.7 mm, optimized for 920 nm, Grintech, Germany), was slowly lowered at an angle of 10-15 degrees (tip towards anterior) such that the prism entered the space between MEC and cerebellum. No aspiration of cerebellar tissue was performed. Once the prism was lowered such that it was visibly below the superficial cortex, it was then pressed forward about 0.5 mm to ensure optimal adherence of its frontal facing surface to the surface of MEC. Similar procedures were used for the implantation of a glass prism (1.3×1.3×1.6 mm, Sunlight, Fuzhou, China), which was glued to a 5 mm diameter cover glass (CS-5R, Warner, MA, USA) and held in place via a custom-made titanium cannula (diameter: 5 mm, 1 mm wall height). Cannula, cover glass and prism were glued together using UV curable adhesive (NOA61, Norland, NJ, USA). The exposed brain around the implantation site was covered with Kwik-Sil (World Precision Instruments) and the clamped lens was carefully fixed to the surrounding bone with UV curable cement (Venus Diamond Flow, Kulzer). A custom-designed titanium headbar was attached and centered on the dorsal surface of the skull and aligned parallel to the top face of the GRIN lens. All exposed areas of the skull, including the headbar, were then covered with dental cement (Paladur, Kulzer), made opaque by adding 0.5 g of carbon powder (Sigma Aldrich, CAS number 7440-44-0). After the surgery, the animals were allowed to recover in a heated chamber (33°C, 30-90 minutes) until they regained complete mobility and alertness.

### Histology

Mice received an overdose of sodium pentobarbital (Apotekforeningen, 100mg/ml) before transcardial perfusion with freshly prepared paraformaldehyde (PFA, CAS Number: 30525-89-4, Alfa Aesar, 4% in phosphate-buffered saline (PBS), P3813, Sigma-Aldrich, Germany). After perfusion, the brain was extracted from the skull and kept in PFA 4% and at 4 degrees Celsius for post fixation. Samples were then sliced on a cryostat (CryoStar NX70, Thermo Scientific, USA, 30-50 μm thick sagittal sections) and either mounted directly on custom made gelatin coated glass slides (for cresyl violet (Nissl) stainings) or collected sequentially in a 24 well plate in PBS (for immunofluorescence). For immunofluorescence stainings slices were incubated for 1 h at room temperature (RT) in blocking buffer (2% gelatine, 2% BSA (bovine serum albumin, Cat#a2153, CAS Number: 9048-46-8, Sigma), 0.1% Triton X-100 (Merck) in PBS). Primary antibodies diluted in blocking solution were added for an overnight incubation at room temperature (RT). After washing with a 1:3 dilution of the blocking buffer in PBS, slices were transferred to a fresh blocking buffer containing fluorophore-conjugated secondary antibodies for 1.5 hours at RT. In some cases, DAPI (1:5000) was mixed into the final washing steps. After a final wash with PBS, slices were mounted in their appropriate anatomical order on glass slides (Polysine adhesion slides (Cat#10219280, Brand: J2800AMNZ, Thermo Scientific, US) using Prolong antifade with DAPI (Cat#P36935, Invitrogen). Primary antibodies used: mouse anti-NeuN, MAB377 (Merck), rabbit anti-GFP, A-11122 (Thermo Fisher). Secondary antibodies used: Alexa Fluor 488, donkey anti-rabbit IgG, A-21206 (Thermo Fisher); Alexa Fluor 647 rabbit anti-mouse IgG, ab169348 (abcam). For confocal imaging, a Zeiss LSM 880 microscope (Carl Zeiss, Germany) was used. Images were then acquired as z stacks using an EC Plan-Neofluar 20×/NA 0.8 air immersion, 40x/NA 1.4 oil immersion (Cat#420762-9900, Zeiss Plan-Achromat, Zeiss, Germany) (Zeiss, laser power: 2-15%).

To determine implant positions and to identify which brain regions were covered by the prisms (Figure S6, see summary across all animals in Figure S2H), histology was aligned to the Allen Mouse Brain Common Coordinate Framework (CCFv3) (Wang et al., 2020) via an open-source python based tool (*brainreg-segment*) that allows for manual annotation of implant positions in a common coordinate space (Tyson et al., 2021) and analysis of brain regions that are covered by the implants.

### 2-photon imaging setup

Custom made two-photon miniscopes were built as described previously (3, 4), but featuring an improved scope body design made out of machinable plastic (PEEK-CF30), which lowered the weight (2.6g) compared to previous versions. The supple fiber bundle (SFB) used in previous versions was substituted by a lighter and more flexible, 0.7 mm diameter fiber bundle with a tapered end piece (TFB), in which scattered fluorescence emerging from the sample is collected over a diameter of 1.5 mm and then coupled into the fiber bundle from the tapered end. A Ti:Sapphire laser (MaiTai Deepsee eHP DS, Spectra-Physics) tuned to a wavelength of 920 nm was used as the excitation source. Pulsed laser light was led through, in series, a half wave plate (HWP, AHWP05M-980, Thorlabs, NJ, USA) in combination with a polarizing beam splitter cube providing power attenuation, an electro-optic modulator (Model 350-80-LA-02, Conoptics, driver: 302RM) providing fast power modulation, and finally three 15-cm long ZF-62 glass tubes (GLA-10×150-AR800-1100, Sunlight, Fuzhou, China), followed by another HWP (AHWP05M-980, Thorlabs). The ZF-62 glass introduces constant positive dispersion, and the laser’s internal dispersion compensation (Mai Tai Deepsee module) minimizes residual dispersion induced by the fiber. The laser light was coupled into a hollow-core photonic-crystal fiber (HC-920-01-FUD, batch 2015, NKT, Denmark) via an aspheric lens (f=11 mm, Geltech, C220TMD-B, Thorlabs) mounted on a fiber launch (MAX350D/M, Thorlabs). The coupling efficiency was monitored with a power meter (S121C connected to PM100D, Thorlabs) and small adjustments were made over time to maintain stable coupling. Coupling efficiency through the HC-PCF averaged around 80-85%.

Inside the scope body the laser was deflected by a MEMS scanner (A3I12.2-1200AL, Mirrorcle, CA, USA), before being led through a scan lens (D0131 / D0166, Domilight, Nanjing, China) and deflected by a dichroic mirror (HT 450-650nm, HR 800-1100nm, Sunlight Ltd., Fuzhou, Fujian, China) towards a custom designed mini objective (water-immersion, D0213, Domilight Ltd., Nanjing, China) and into the sample. Fluorescent light from the sample was led through the TFB into the detection module, consisting of an aspheric condenser lens (ACL25416U-A, Thorlabs) for collimation, low-pass filter (HC 720/SP, Semrock, AHF Analysentechnik AG Tuebingen, Germany), a dichroic beamsplitter plate (HC BS 560, Semrock, AHF Analysentechnik AG), followed by two transmission filters (“green channel”: 525/50 BrightLine HC, Semrock; “red channel”: 620/60 ET Bandpass, Chroma; AHF Analysentechnik AG), and two aspheric condenser lenses in each detection path (ACL25416U-A, Thorlabs) focusing the light on two GaAsP photomultiplier tubes (PMT2101, Thorlabs). A mechanical shutter (SHB1, Thorlabs) was put in front of the detection module to protect the PMTs from unexpected exposure. The MEMS driver (BDQ PicoAmp 5.4 T180, Mirrorcle, CA, USA) was mounted in a custom-made housing and connected to the MEMS scanner via a bundle of thin isolated wires.

The miniscope control stack consisted of NI hardware (NI PXIe-7961R NI FlexRIO FPGA Module, 3x PXIe-6341, X Series DAQ, NI 5734 Digitizer, chassis: NI PXIe-1073, BNC breakout boards: BNC-2090A) controlled via scanimage (Pologruto et al., 2003) (v.5.5, Vidrio Technologies, Virginia, USA). The MEMS scanner was driven in “resonant mode” in scanimage at 2 kHz resulting in a framerate of ∼7.52 fps at 512×512 pixels. The average laser power before entering the GRIN lens was usually below 150 mW.

### Miniscope imaging

At the beginning of every imaging session and for general checks of GCaMP expression, mice were restrained through their headbar, with their limbs resting on a freely rotating treadmill consisting of a ∼85 by 70 mm (radius x width) styrofoam wheel with a metal shaft fixed through the center. Low friction ball bearings (HK 0608, Kulelager AS, Molde, NO) were affixed to the ends of the metal shaft and held in place on the optical table using a custom mount (https://github.com/kavli-ntnu/wheel_tracker). A table-top fluorescence microscope (SFM, Thorlabs), equipped with a 470 nm excitation source (M470L3, Thorlabs) and “GFP” excitation / emission / dichroic filter set (MDF-GFP2, Thorlabs) was used both to acquire epifluorescent images of chronic implants and to maneuver the two-photon miniscope over the mouse skull via a custom-made holder, which itself attached to the objective of the table-top microscope (MY5X-802-5X, Mitutoyo). The miniscope itself was held in place on the animal’s head via a custom holder (“cup”), which was cemented in place above the implanted GRIN lens (with dental cement, see “GRIN / prism implantation” above), while imaging with the miniscope to determine the most suitable medio-lateral and anterior-posterior (=dorso-ventral through GRIN+prism) position. This procedure yielded a field-of-view (FOV) that was ideally both centered on MEC and had a high number of visible cells. For every subsequent recording the miniscope was inserted into the cemented cup in the same manner as during cementing / fixation, which enabled reliable re-finding of the FOV of previous recording sessions. The miniscope was then secured in place via two M1.6 set screws offset at 90 degrees in the holder base. After releasing the scope from its holder on the table-top microscope, two RGB LEDs (NeoPixel Nano, PID 3484, Adafruit, New York, USA) were mounted to the scope body on a light plexiglass bar (length: ∼43 mm), which allowed tracking of both the animal’s position in x and y and its head rotation in open field arenas.

Mouse behavior in open field arenas was recorded in RGB via a camera mounted to the ceiling of the recording room (acA2040-55uc, Basler, Ahrensburg, Germany; objective: AZURE-1236ZM, RMA Electronics. Inc.). Brightness of the RGB LEDs (one set to only emit blue light, the other to only emit red light) was adjusted to enable smooth and reliable tracking via a custom written application that allowed both online tracking and visualisation of animals’ paths as well as parallel saving of recorded videos and extracted position data at 500×500 px and 40 Hz (https://github.com/kavli-ntnu/2P_tracking). Imaging and tracking were synchronised via TTLs (Frame clock of two-photon miniscope and exposure of new tracking frames (Basler camera GPIO interface)), which were recorded via WaveSurfer, an open-source software package for acquiring neurophysiology data in Matlab (https://wavesurfer.janelia.org/), on a NI PCIe-6321 card at 5 kHz. WaveSurfer triggered the miniscope acquisition and generated trigger pulses at regular intervals (40 Hz) for the tracking camera, which in turn sent TTLs for every exposure back to WaveSurfer.

### Analysis pipeline

All analyses, from extraction of transients, deconvolution of fluorescence data (see below) to the analysis of spatial tuning and topographic properties were integrated in a custom pipeline using the Datajoint framework (https://datajoint.io/), an open-source library for programming scientific databases and computational data pipelines (Yatsenko et al., 2015).

### Analysis of imaging time series

Two-channel imaging timeseries recorded via ScanImage (Pologruto et al., 2003) were analyzed using the *Suite2p* python library (https://github.com/MouseLand/suite2p), an open-source analysis pipeline for processing of two-photon calcium imaging data (Pachitariu et al., 2017). We used its built-in motion correction, region of interest (ROI) extraction and spike deconvolution routines. For tdTomato cell analysis in Figure S3D, *cellpose* (https://github.com/MouseLand/cellpose), an open-source generalist algorithm for cell and nucleus segmentation (Stringer et al., 2020) was used on second channel average projections that were corrected by subtracting a linear offset extracted by regressing first against second channel projections for each session.

For calcium time series data, non-rigid motion correction was chosen to align each frame iteratively to a template and visual inspection of the corrected stacks and built-in motion correction metrics were used to judge its quality. After automatic ROI extraction, the *Suite2p* GUI was used to manually sub-select putative neurons based on anatomical and signal characteristics and discard obvious artefacts that accumulated during the analysis (e.g., unrealistically large ROIs or ROIs that did not have a clearly delineated footprint in the generated maximum intensity projection). Raw fluorescence traces were processed to create a corrected fluorescence signal (F_corr_) by first subtracting 0.7 * neuropil signal, and ΔF/F was then calculated as (F_corr_ -F_0_) / F_0_. F_0_ was estimated as constant for each cell after smoothing F_corr_ with a 15 s spanning window (convolution of a scaled window with the signal, after introducing reflected copies of the signal at both ends so that transient parts are minimized in the beginning and end part of the output signal) and taking the median of the minimum 20% of this signal. Non-negative deconvolution (Friedrich et al., 2017) was used for deconvolving the neuropil and baseline corrected fluorescence (F_corr_), yielding unfiltered, deconvolved events. After deconvolution we kept events (i.e., filtered events) that were larger than one standard deviation over the mean (calculated over all extracted events for that neuron) and filtered out cells that did not meet a signal-to-noise ratio (SNR) cutoff of 3.5. We used the deconvolved, filtered amplitudes (events) as input for all subsequent analyses (spatial tuning maps etc.).

For SNR calculation, noise statistics were extracted from the ΔF/F traces after filtering for episodes in which no deconvolved events were present (F_noise_). These episodes had to lie at least 1 second before and 10 seconds after any deconvolved event to prevent signal contamination (safety margins). The SNR of that cell was then calculated as the ratio of the mean amplitude of filtered events over the standard deviation of F_noise_. If no filtered events were maintained for a cell, a SNR of zero was assigned. Since the MEMS scanner of the two-photon miniscope introduces warping artefacts to the image due to inhomogeneities in its control voltage response, unwarping procedures were used to re-align ROI pixel and image projection data (i.e., average or maximum intensity projections). For this, a standard grid distortion target (R1L3S3P, Thorlabs) was first imaged and piecewise affine transformation was used to re-align warped key points (crossings of all grid lines) to an idealized grid pattern. The resultant transformation matrix was then applied to unwarp both projection data (e.g., average, maximum-intensity projections) as well as ROI data. We filtered out ROIs that retained less than 10% of their original number of pixels after unwarping to get rid of edge artefacts for topographic analyses.

Since the GRIN lenses we used have a magnification of 0.8 between the object plane and image plane), anatomical data that was acquired via these implants underwent additional post-correction with a scaling factor that was extract by aligning 2p miniscope FOVs to images acquired on the table-top fluorescence microscope through the same implant (see hardware description above, example can be seen in Figure S1C bottom).

### Analysis of spatial tuning properties

Tracking (position of 2 LEDs on the mouse head) and imaging data (deconvolved, filtered fluorescence for each ROI) were synchronized by timestamp. As the tracking sample rate was ∼5 times greater than that of imaging, each ROI could be assigned tracking data with greater temporal precision by assigning a small temporal offset to the ROI data between the start and end time of each imaging frame acquisition (∼133 ms per frame), depending on the time it took the laser to reach that ROI within each FOV. Raw position and angular (head direction) data was smoothed with a Gaussian kernel (sigma 50 ms) and the instantaneous speed of the animal was then calculated. We applied speed cutoffs after smoothing the speed signal with a Gaussian kernel (sigma 150 ms, lower limit: 25 mm/s, upper limit: 500 mm/s) before analyzing any spatial tuning. Spatial data was binned (25 × 25 mm) and smoothed with a Gaussian kernel (sigma 50 mm). Spatial tuning maps were created by dividing the binned imaging data (filtered, deconvolved fluorescence) by occupancy in each bin. Similarly, for head direction tuning analyses, the angular signal was smoothed with a Gaussian kernel both temporally and spatially (bin size 2 degrees, temporal smoothing sigma 4 degrees, spatial smoothing sigma 6 degrees). The open-source package *opexebo* (https://pypi.org/project/opexebo/) was used to analyze all tuning properties from spatial tuning maps and angular tuning curves. Perceptually uniform sequential colormaps from the *CMasher* python library (van der Velden, 2020) (v1.6.1) were used for display of 2D tuning maps.

### Field detection in spatial tuning maps (*opexebo*)

Place fields in spatial tuning maps and 2D autocorrelations (see Grid Score calculation) were identified by using an adaptive threshold method. First, local maxima (peaks) were found using *SEP* (Barbary, 2016) (python library for source extraction and photometry, https://pypi.org/project/sep/). Then, looping over every single peak, an initial threshold was determined (usually 0.8 of maximum) that, when lowering it in small increments, would yield an expansion of the field area around that peak. This field specific starting threshold was then systematically lowered in steps of 0.02 while field properties were measured until it reached one of several stopping criteria: 1. the field area could not be determined, 2. voids (“holes”) appeared within field, 3. the expansion of the field lead to inclusion of other field peaks, 4. the field size more than doubled in one step or increased by the same or greater proportion in three consecutive steps, or 5. the field size did not change over 10 consecutive steps. Finally, fields were filtered out that had mean amplitudes below 10% of the global maximum or spanned fewer than 5 connected bins in the spatial tuning map.

### Grid Score calculation (*opexebo*)

Spatial autocorrelograms of spatial tuning maps were used to obtain a measure of 60-degree place field periodicity (grid score). Peaks in spatial autocorrelations were found as described above (“Field detection in spatial tuning maps”). The grid score was calculated by expanding a circle around the center field and calculating a correlation value of that circle with rotated versions of itself. For each radius the score was calculated as the difference of the minimum correlation value at 60 and 120 and the maximum value at 30, 90, and 150 degrees. The final gridness score value was then obtained by calculating a maximum over a sliding mean of expanding circles. To determine grid spacing (distance of fields to center peak) and orientation, the orientation and distance of the six closest fields to the center peak was quantified. If two fields had less than 20 degrees difference from each other, the one with the larger distance from center was discarded. The first three fields were maintained, starting from the horizontal axis, and progressing in ascending (angular) order. The average orientation was extracted as the average difference to 60 degrees axes.

### HD tuning curves and mean vector length

For circular statistics, we adapted scripts from the *circstat* toolbox (Berens, 2009) To obtain head direction tuning curves, filtered and deconvolved fluorescence was binned in 2 degree radial bins and divided by occupancy. Tuning curves were smoothed with a Gaussian kernel (sigma 6 degrees). The mean resultant vector length (mean vector length, MVL) for circular data was then calculated as described previously (Rowland et al., 2018) and ranged from 0 to 1, where 1 is the case that all data are concentrated in the same direction. The calculation is described in (Zar, 2010), Section 26.4.

### Border score (*opexebo*) and boundary vector score

The border score was calculated for spatial tuning maps as in (Solstad et al., 2008). This score is in the range [-1, 1] and reflects both the width of a field (what fraction of a single wall it touches), and the depth of a field (how far away from the wall it extends). It was only evaluated for the single firing field that had the greatest wall coverage. All other fields were ignored. The highest scores are returned for cells that have a field both with maximum width along one wall of the square open field and shallow depth. Scores are usually below ∼0.9 in a typical spatial tuning map (limited by the width of the binning compared to the arena extent). To account for additional firing fields, fields that had elongated fields parallel but offset from one of the walls, and those cases in which fields were less perfect (i.e., shallow) but still visibly border-like we calculated a boundary vector score (BVS). This score is in the range 0 to 1, where 1 is assigned to cells that have perfect and isolated border fields, which may or may not be offset from one of the walls. For this, the spatial tuning map of a cell was first binarized, with bins above median + 1 standard deviation over all bins set to one and zero for the rest. Then connected regions (=fields) were detected and quantified and those that covered fewer than 16 bins or had maximum amplitudes below the median + 2 standard deviations over all bins in the unprocessed spatial tuning map were discarded, thus yielding a filtered field map. Bars of variable thickness (from 1 to 5 bins) spanning the whole width / height of the spatial tuning map were then moved over the map in x / y and at each position a score was calculated as: Overlap_Bar_ -r * Overlap_Rest_, where Overlap_Bar_ is the fraction of bins in the bar overlapping with a field and Overlap_Rest_ is the fraction of bins overlapping with a field outside the bar. The r-value is a factor in range 0 to 1, which weighs the contribution of extra-bar field bins in decreasing the score and was set to 0.5 in this study. The maximum score over all different combinations of bar widths and position offsets was chosen as the boundary vector score for that cell. The BVS was preferred over the border score when a more graduated evaluation of border responses was needed (i.e., topographic tuning maps).

### Shuffling distribution and cutoffs for spatial scores

Shuffled distributions of spatial scores were created for each cell by randomly shifting the tracking versus the deconvolved fluorescence signal in time. Data points that were shifted beyond the last element were reintroduced at the first. A total of 500 shuffle iterations were introduced per cell, with possible shifts being at least 0.5 seconds apart to account for the fact that due to slow time courses of the calcium sensor, neighboring timepoints could not be considered independent samples. A safety margin of 2 seconds was introduced at both ends of the timeline. Spatial scores were calculated for each shuffling iteration and each cell’s real score was compared to percentile cutoffs of a session wide distribution (shuffling results over all filtered cells per session). Shuffling cutoffs used throughout the study were the 95^th^ percentile of shuffled distributions unless otherwise indicated.

### Object vector cells

Object vector cells (OVCs) were usually identified over three adjacent sessions (15-20 minutes each), comprising a base session without objects and two object sessions. For this three-session design, one object (a 64×64×225 mm colorful duplo brick tower) was placed in the middle of one of the open field quadrants in the first object session and moved to the middle of the diagonally opposite quadrant in the second object session. In rare cases only two sessions were run (one base and one object session) and similar looking objects were placed in opposing quadrants of the open field. To analyze object related firing patterns, object centered occupancy and tuning maps (object vector maps) were first created as described previously (Høydal et al., 2019), by tiling space around the center of each object in 72 radial wedges, which were further subdivided along distance from the center into 2.5 cm wide bins. The object vector score (OV score) was then defined as the 2D correlation between the two object vector maps. Fields in the spatial tuning map were analyzed as described above (“Field detection in spatial tuning maps”) and for every field, an object centered distance and angle was extracted, from the center of the object location to its centroid. Then, putatively matching fields in the spatial tuning map across objects (either two sessions with one object each or two objects in one session) were identified by calculating all pairwise Euclidean field distances in object centered coordinates and keeping the one with minimum distance to each other. To filter for OVCs, a range of cascading criteria were used: 1. OV scores had to be above the 95^th^ percentile shuffling cutoff (evaluated on cellular, not session level), 2. Spatial tuning maps had to have information content above the 95^th^ percentile shuffling cutoff in each of the object sessions (evaluated on cellular, not session level). Information content was calculated as described in (Skaggs et al., 1993) ([bits / deconvolved fluorescence]), 3. The maximum distance in object centered space between putatively matching fields in both object sessions had to be below 250 mm and they were required to have a minimum distance to the object of 40 mm, and finally 4. fields had to show at least 50% increase in activity compared to the same location in the baseline session. These criteria were used to ensure that putative OVCs developed clearly delineated firing fields in object sessions (information content and ratio of activity to base sessions) and that those fields were in a distance / angle relationship that matched expectations for OVCs (ensured by both OV score and distance cutoffs).

### Grid module extraction

Example data shown in Figure S6D. We ran principal component analysis (PCA) on arrays of spacing versus orientation data from a subset of grid cells with high grid scores (>99^th^ percentile shuffling distribution cutoff) in each animal. After dimensionality reduction, unsupervised clustering (*HDBSCAN*, https://github.com/scikit-learn-contrib/hdbscan) was run on this dataset of to extract seed clusters (i.e., preliminary grid modules; “skeleton”, Figure S6D top). A k-nearest neighbor classifier (kNN implementation in scikit-learn (Pedregosa et al., 2011), v0.24.2) was then trained on this data; this trained classifier was subsequently applied to expand each module by adding grid cells with >95^th^ (and <99^th^) percentile shuffling cutoff (Figure S6D bottom). Grid modules that contained fewer than 5 cells were discarded.

### Session FOV alignment

Due to changes in the exact clamping angle and position of the miniscope in its holder (“cup”) from day to day, the field of view (FOV) had to be aligned to a reference before projections over multiple recordings could be created (described in detail below). The goal was to create alignments on the coarse macroscopic level, maintaining alignment of for example the blood vessel pattern and not necessarily on the (sub-)cellular level. Matching anatomical landmarks were picked in both the GCaMP (“green”) and tdTomato (“red”) channels for every pair of reference / session FOVs. The reference FOV was first picked, which showed maximum anatomical overlap with most other FOVs. Then landmarks were picked in a custom written software (*PyQT5* GUI), which allowed inspection of both cellular details (average and maximum intensity projections) and blood vessel patterns (average intensity projections) in both channels. At least three pairs of points had to be defined for successful alignment (number of point pairs: (mean ± SD) 7±2). The reference was first padded and nudged to create sufficient space for the creation of a stitched composite. FOVs were then warped into a common coordinate space through an affine transformation defined over the user-defined point pairs. To evaluate the increase in anatomical overlap after versus before alignment of recorded FOVs, the structural similarity index (SSIM) (Wang et al., 2004) was calculated for same-sized areas in the average projections of aligned and reference session FOV, and compared before and after alignment. An increase in SSIM signified that the alignment yielded an improvement in anatomical overlap. The average projection was chosen because it maintained anatomical information (e.g., blood vessel patterns) while blurring cellular details that would otherwise disturb the alignment assessment.

### FOV score map projections, Moran’s I and correlations

To obtain FOV score maps (Figure 6 and Figure S6C), anatomical composites were first created as described above (“Session FOV alignment”) and the boundaries of each composite were used to create binning vectors (bin size 5 μm). Unwarped and aligned ROI pixel data was then binned in x and y (filtered session ROIs). Each 25 μm^2^ bin was assigned a score defined as the maximum over all pixels inside that bin (e.g., max. grid score of all ROI pixels in each bin). Bins that did not contain any data in the first place were set to NaN. Since most spatial scores tend to not fall off smoothly at the lower end of their range (for example grid scores or border scores), pixels containing values below shuffling cutoff were set to zero. Unless otherwise indicated, the 95^th^ percentile shuffling cutoff (session level) was used for grid, border and HD (MVL) scores and combined criteria as described above were used for object vector cells. The final FOV score maps were then obtained by taking the average projection of the aligned and binned score data of each session FOV. For border cells, the boundary vector score is plotted, for reasons indicated above, i.e., to obtain smoothness of values across the possible range of scores. To analyze spatial clustering in score maps, Moran’s I, a global autocorrelation statistic, was used (Cho, 1983). It is a measure of how similar neighboring bins in a 2D map are to each other and captures the intuition that spatial clustering introduces dependencies of value changes in bins over space compared to a “salt and pepper” organization in which no such dependencies can be observed. Moran’s I values were calculated via the *ESDA* library, an open-source python library for the exploratory analysis of spatial data (sub module of *PySAL*, Rey and Anselin, 2010). To obtain a significance level and compare each Moran’s I value to a “salt and pepper” organization, shuffled score maps were created by permuting cell IDs across all cells for all sessions that were used in the original projection. Random samples of shuffled maps for each of the sessions in the original projection were then combined and Moran’s I values were calculated as for the real data. This process was repeated 5000 times to create a distribution of Moran’s I values and extract 1^st^/5^th^ and 95^th^/99^th^ percentile cutoffs respectively. FOV score map projections with Moran’s I values that were above the 95^th^ percentile cutoff are called “clustered” since they deviate most strongly from a salt and pepper (i.e. random) organization obtainable from the same data.

To determine the relationship of cell class territories over anatomical space, FOV score maps were first smoothed with a Gaussian (sigma 10 microns) and then the 2D correlation was calculated. The correlation value obtained from this comparison was compared to a shuffled distribution obtained by correlating randomized FOV score maps to each other. The procedure for obtaining randomized maps was identical to the one described above for Moran’s I values. 10,000 comparisons were drawn, and 1^st^/5^th^ and 95^th^/99^th^ percentile cutoffs were determined from the resulting distributions. Correlation values below the 5^th^ percentile cutoff were considered “anti-correlated” and those above the 95^th^ percentile correlated. Perceptually uniform sequential colormaps from the *CMasher* python library (van der Velden, 2020) (v1.6.1) were used throughout.

### Masking MEC and adjacent structures

In some mice that were implanted more medially than average and that were labeled with retrogradely transported AAVretro-CAG-tdTomato in hippocampus, a clear drop-off of tdTomato fluorescence could be observed in parts of the FOV, demarcating the boundary between MEC (high expression of tdTomato) and adjacent structures (i.e., parasubiculum PAS, low expression). We used a custom-written workflow built in napari (https://napari.org/) to paint in anatomical masks for MEC and adjacent structures. Where no tdTomato boundary was observable, we relied instead on Nissl-stained histology to infer the putative FOV position and subsequently the anatomical region that was most likely covered by the implant. Extracted masks were used to filter ROI data and label each cell by its anatomical region.

### Nearest neighbor (NN) analysis

Intra-cell class nearest neighbor distances (Figure 4, S4, 5 and S5) were calculated with the help of *PySAL* (Rey and Anselin, 2010) (https://pysal.org/, v2.3.0) and pairwise distances and inter-cell class distances (KDTree for fast generalized N-point problems) with tools from *scikit-learn* (Pedregosa et al., 2011) (https://scikit-learn.org/, v0.24.2). Unless otherwise indicated, shuffled distributions were created over n=1000 iterations and the median over all shuffled values was used for any normalizations involving results of this shuffling. For inter-cell class distances (i.e., two populations A and B, Figure 4 and Figure S4) size matched groups were first created by randomly picking groups of cells that had the same number as the smaller population (A or B) in each case. For example, if population A contained 50 cells above threshold (starter cells), and population B 25 cells, A was subsampled to only contain 25 cells. This picking procedure was repeated 1000 times and the median over all permutation results was used for further calculations. Then, two population statistics could be extracted, i.e., reaching from class A to class B or class B to class A. In the idealized case that half of the FOV was covered by one or the other cell population, the directionality did not have an effect. However, because of FOV edge effects, differences could be quite pronounced. To not get biased towards either side, we took the average value of over those two statistics.

### Graph analysis and spring-loaded model

For graph analysis in Figure 4E we used an implementation of an undirected graph that can hold multiple nodes from the open source *networkx* python library (https://networkx.org/, v2.5). Weights between nodes (different cell classes) were assigned by the inverse of the mean over the nearest neighbour distances between two cell classes (see also “Nearest neighbour (NN) analysis” above) and were centered around zero. To introduce (random) variability before running the simulation, node positions were randomly initialized in a square centered on (0,0). The simulation itself utilizes the *networkx* implementation of the Fruchterman-Reingold algorithm (*networkx*.*drawing*.*layout*.*spring_layout*). This algorithm simulates a force-directed representation of the network, treating edges as springs holding nodes close, while trying to reach an equilibrium state at every step at the simulation by re-arranging node positions according to the weight of their edges. The simulation was run for 1000 steps for Figure 4E and Figure S4D, but varied systematically in Figure S4E. To quantify the divergence of the “Grid” (grid cell) node compared to all other nodes, compensating for the varying distances that could occur in the simulation, we calculated the distance ratio of grid to all other cell nodes over the mutual distance of all other cell nodes (i.e., border, HD and OV nodes). The distance of grid to all other nodes was quantified as the Euclidean distance between the grid cell node and the center of all other nodes, and the mutual distance of all other (non-grid) nodes as the mean of pairwise distances between those nodes. Ratio values close to 1 indicate that the divergence between grid to all other nodes was similar to the distances in between all other (non-grid) nodes.

## QUANTIFICATION AND STATISTICAL ANALYSIS

The details of statistical testing, including the test statistic, p-values, number of results and what the number of results refer to can be found in the figure legends throughout. Definition of distribution centers and dispersion for each figure are indicated in the figure legends and described in the main text. Randomization procedures are described in detail in the associated sections in the *Method details* section and explained in the main text and figure legends where appropriate. Most statistical testing was performed using non-parametric tests because Gaussian sample distributions could not be ascertained due to low sample numbers. All test statistics and p-values were calculated using *SciPy*’s stats module (v1.5.4) (Virtanen et al., 2020) and a significance level of 0.05 was used throughout to assess whether results were statistically significant (p < 0.05) or not (p ≥ 0.05).

