## Supplementary figures and legends for "Functional network topography of the medial entorhinal cortex"

**This file includes:**

All supplementary figures and associated legends

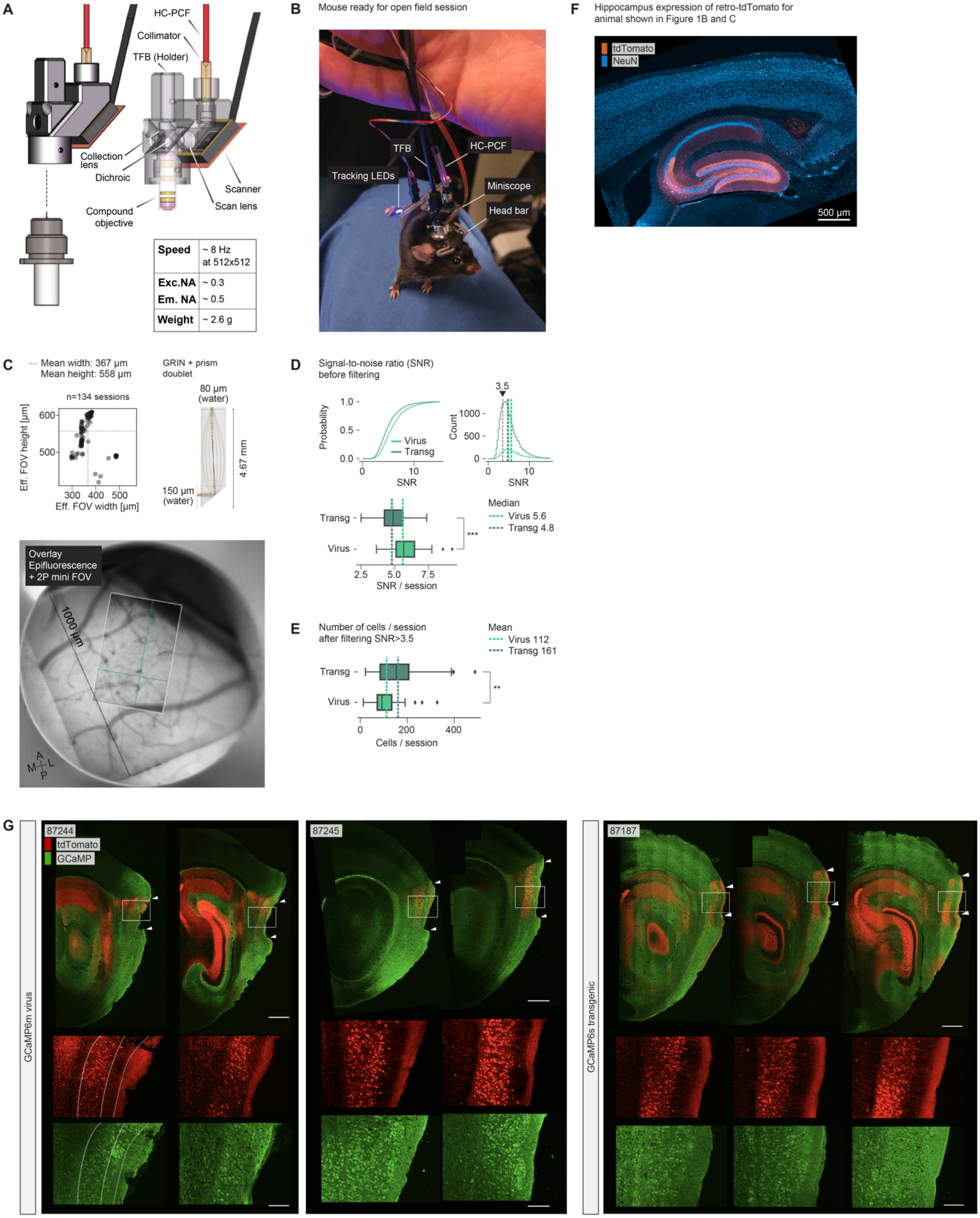
A
HC-PCF
Collimator
TFB (Holder)
Collection lens
Dichroic
Scanner
Scan lens
Compound objective
Speed ~ 8 Hz at 512x512
Exc.NA ~ 0.3
Em. NA ~ 0.5
Weight ~ 2.6 g
B
Mouse ready for open field session
TFB
HC-PCF
Tracking LEDs
Miniscope
Head bar
F
Hippocampus expression of retro-tdTomato for animal shown in Figure 1B and C
tdTomato
NeuN
500 µm
C
Mean width: 367 µm
Mean height: 558 µm
n=134 sessions
Eff. FOV height [µm]
Eff. FOV width [µm]
GRIN + prism doublet
80 µm (water)
150 µm (water)
4.67 mm
Overlay Epifluorescence + 2P mini FOV
1000 µm
A M L P
D
Signal-to-noise ratio (SNR) before filtering
Probability
Virus
Transg
SNR
Count
3.5
Median
Virus 5.6
Transg 4.8
***
SNR / session
E
Number of cells / session after filtering SNR>3.5
Mean
Virus 112
Transg 161
**
Cells / session
G
87244
tdTomato
GCaMP
87245
87187
GCaMP6m virus
GCaMP6s transgenic

**Figure S1. 2P miniscope, field of view (FOV), signal characteristics and example histology. Related to Figure 1.**
**(A)** Schematic of the 2P miniscope. Table shows characteristics of the scanner and objective combination used in this study. The light scope body and flexible connection cable are as described for the miniscope version in the accompanying Resource article by Zong et al. ("MINI2P"). However, this version of the miniscope lacks the built-in Z-focusing module and the ability to shift FOVs systematically between sessions via a stitching adapter. HC-PCF: Hollow-core photonic crystal fiber, TFB: tapered fiber bundle.
**(B)** Photo of a mouse carrying the 2P miniscope with annotations (TFB: Tapered fiber bundle (signal detection); HC-PCF: hollow-core photonic crystal fiber). The weight of all components mounted on the animal head was counterbalanced by a pulley system while animals were running inside the open field arena.
**(C)** Left: Effective (Eff.) FOV height and width (after motion correction) of 134 recorded sessions. Stippled blue lines indicate the mean (Mean width ± SD [µm]: 367 ± 40, Mean height ± SD [µm]: 558 ± 50 µm).
Right: Schematic of custom GRIN+Prism implant and its optical properties (Image side working distance: 80 µm in water; object side working distance 150 µm in water). Bottom: Overlay of two-photon imaging FOV (rectangle in center) and epifluorescence image (whole image) through the same implant *in vivo*.
**(D)** Comparison of signal-to-noise ratio (SNR) across GCaMP6 transgenic ("Transg") and virus ("Virus") injected animals; n animals: 12 transgenic; 3 virus. Top: Distribution plots (left and right) across all cells. Colored, stippled lines indicate median; grey stippled line indicates SNR cutoff (3.5) used throughout this study to filter cells. Bottom: SNR per session. Box shows range from first to third quartile, whiskers indicate first / third quartile ± 1.5 times the inter quartile range (IQR), thick vertical line indicates median. The median SNR of cells recorded in transgenic animals was slightly lower than in virus injected animals (Median: Transg 4.80, Virus 5.61; Mann-Whitney U=844, $n_{Transg}$=103, $n_{Virus}$=31, p=0.000073***, two-sided).
**(E)** Comparison of number of cells per session across GCaMP6 transgenic ("Transg") and virus ("Virus") injected animals after filtering; n animals: 12 transgenic; 3 virus (Mean±SD [n cells]: Transg 160.7±96, Virus 111.5±68; Mann-Whitney U=1071, $n_{Transg}$=102, $n_{Virus}$=31, p=0.0033** two-sided). Box plot characteristics as in (D).
**(F)** Confocal image of a stained, sagittal brain slice of the animal shown in Figure 1 B and C, which shows expression of AAVretro-CAG-tdTomato in DG and CA3 of hippocampus, close to the injection site. Red: tdTomato, native fluorescence; blue: primary mouse aNeuN + 647 nm secondary.
**(G)** Example sagittal slices of 3 animals with high fractions of functional cell types. Insets at the bottom of each image are magnifications of the white boxed regions. Green: GCaMP6 (aGFP), red: retro-tdTomato (native fluorescence). GCaMP virus injected animals: 87244, 87245; GCaMP transgenic animal: 87187. Dense cellular labeling was observed in the superficial layers of MEC in all animals. White arrow tips indicate the dorsal end of MEC (top) and most ventral imprint of prism (bottom). Depression of the brain surface surrounding the white box reflects the location of the prism. An example layer annotation (Layer 1 to 3) is shown for animal 87244 (left). Scale bars 500 µm, insets: 125 µm. Additional pictures of GCaMP and tdTomato expression in sagittal brain slices are shown for animal 88592 (GCaMP transgenic animal) in Figure 1C and Figure S3D and animal 88106 (GCaMP transgenic animal) in Figure S3D.

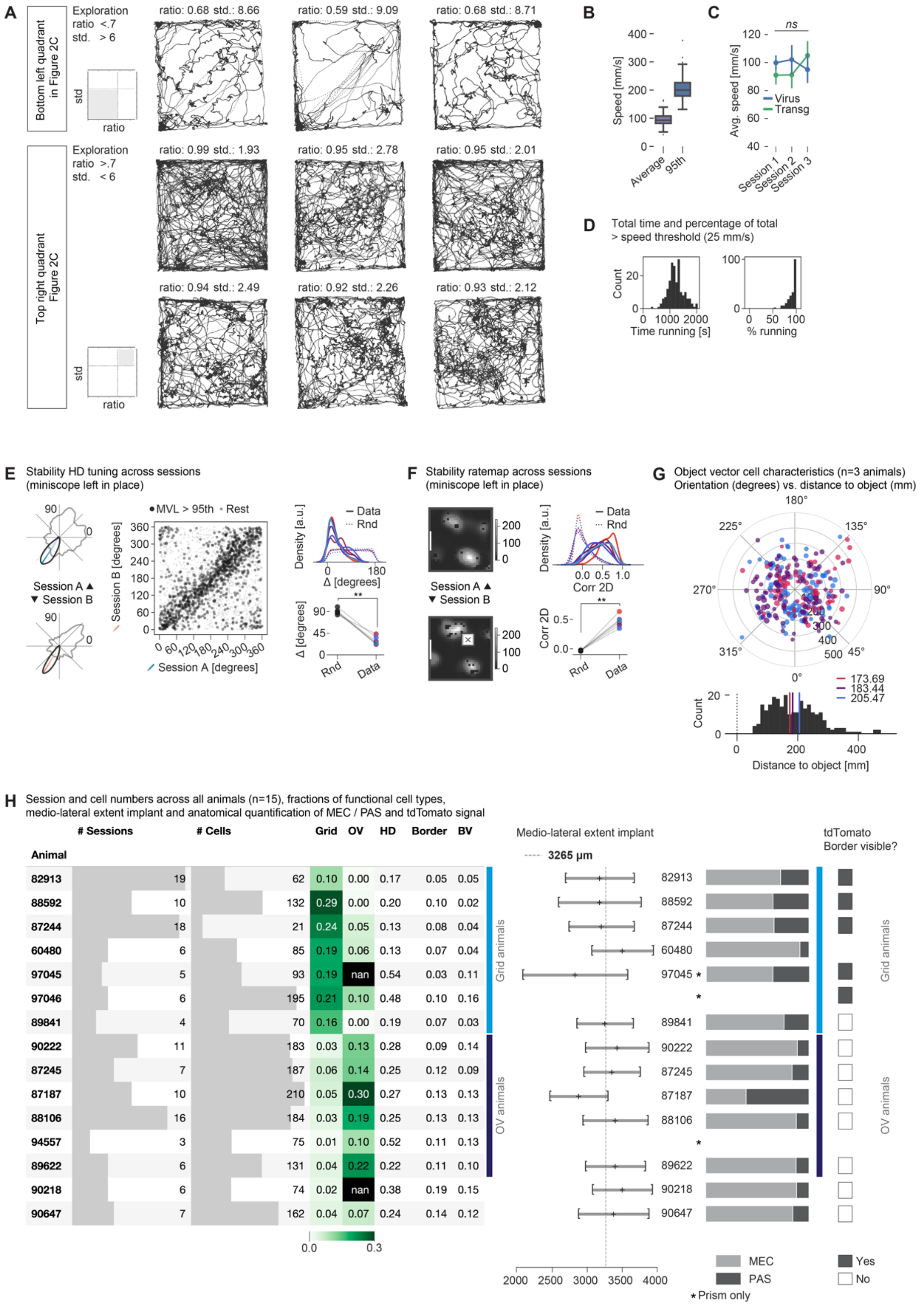


**Figure S2. Behavior, tuning stability, OV cell characteristics and summary of recorded cells and implant locations. Related to Figure 2, 3 and 6.**

**(A)** Example path plots of sessions in the bottom left (top) and upper right (bottom) quadrants in Figure 2C. Exploration ratio (“ratio”) and standard deviation (“std”) values are shown above each plot.
**(B)** Speed comparisons (same data as analyzed for Figure 2C). Left: Average and 95$^{th}$ percentile (95$^{th}$) of speed distributions across recording sessions; Average speed ± SD: 94.7 mm/s ± 20.6; 95$^{th}$ percentile speed ± SD: 203.7 mm/s ± 37.5.
**(C)** Speed across multiple, consecutively run sessions (<60 seconds break between adjacent sessions) shows that animal behavior stayed constant (e.g., across an open field and two object sessions) (figure shows mean ± 99$^{th}$ CI; Kruskal-Wallis H=3.68, $n_{Session1}$=123, $n_{Session2}$=48, $n_{Session3}$=32, p=0.159 *ns*); ns, not significant (p>0.05); virus injected WT animals: blue; transgenic GCaMP animals: green.
**(D)** Distribution of time spent running, i.e., above the speed threshold of 25 mm/s (left, absolute; right, relative to whole session length) for recordings shown in Figure 2C; time: 1304.8 s ± 479.8 (mean ± SD); percentage of whole session spent running: 91.5% ± 8.8 (mean ± SD).
**(E+F)** Stability of single cell head direction (HD) tuning curves and spatial tuning maps across sessions. Tuning stability from one session to the next (session “A” and “B”; miniscope not detached between sessions, the animal is sometimes taken out of the box, handled, and put back in).
**(E)** Tuning stability of head direction cells recorded in two sessions. Left: Tuning curves of one cell recorded in two consecutive open field sessions (“A” and “B”). Blue and orange lines show (approximate) average angle of the tuning curve. Scatter plot in the middle shows average tuning angle in Session A vs. B for 2,615 cells across 6 animals with thick black dots indicating cells that met the 95$^{th}$ percentile shuffling cutoff (1,112 cells > MVL 95$^{th}$). Dots correspond to individual cells. Right: Absolute difference in degrees of tuning angle in session A vs. session B (Data, colors indicate animals, n=6), compared to randomly (Rnd) picked cell pairs. All cells filtered by MVL > 95$^{th}$ percentile. Top: Kernel density estimate, bottom: Median per animal. The median tuning angle difference is significantly lower for analyzed cell pairs, compared to random, co-recorded cell pairs (Mann-Whitney U=0, $n_{Data}$=6, $n_{Rnd}$=6, p=0.0025**, two-sided). Note that the Gaussian kernel used in the density estimate (top) introduces artificial smoothing at the edges of the data range (0 and 180 degrees).
**(F)** Tuning stability of cells with high information content (>95$^{th}$ shuffling cutoff for information content). Left: Spatial tuning maps of one cell recorded in two adjacent sessions (session “B” is an object session, object location indicated by cross in white box). Analysis as in (E), but with spatial tuning map correlation values instead of tuning angle difference (Mann-Whitney U=0, $n_{Data}$=6, $n_{Rnd}$=6, p=0.0025**, two-sided).
**(G)** Distribution of field orientation and distances to objects in OV cells match previously published results with tetrodes (colors=animals, n=3 animals, 260 cells). Bottom: Histogram of field-to-object distances: dashed line indicates 0, vertical-colored lines indicate the mean distance to objects over all fields for each animal.
**(H)** Left: Number of sessions (# Sessions) and mean number of cells (# Cells, after filtering) per animal. Size of the grey bar in the background scales with the absolute value in each table entry. Next, the mean fraction (compared to all filtered cells in a recording) of grid, object vector (OV), head direction (HD), border and boundary vector (BV) cells per session (> 95$^{th}$ percentile shuffling cutoff) are shown. Grid and OV fractions are color coded (colorbar is shown at the bottom). Animals are ordered by “Grid animals” (blue bar, ≥ 10% grid cells on average), “OV animals” (violet bar, ≥ 10% OV cells on average) etc. Middle: Implant positions across the 13 animals analyzed in this study. Sagittal, Nissl-stained brain slices were aligned to the Allen mouse brain common coordinate framework (CCFv3) and implant covered regions were manually annotated. The line chart in the middle shows the medial to lateral extent of the implant, with the ‘+’ denoting the midpoint. The dashed vertical line (light grey) indicates the average across animals (~3.3 mm lateral from midline). The horizontal bar plots towards the right indicate the approximate ratio between the volume covered by MEC (light grey) vs. PAS (dark grey). Grid animals were on average implanted more medially compared to OV animals. Right: Indication of whether a clear border was visible in the retro-tdTomato signal - indicating the transition between MEC (lateral) and PAS (medial) - either at the single session level or after stitching all session FOVs together (dark grey - yes, empty box - no). Where no box is shown, no retro-tdTomato was injected. This medio-lateral region boundary was visible in most grid animals injected with retro-tdTomato.

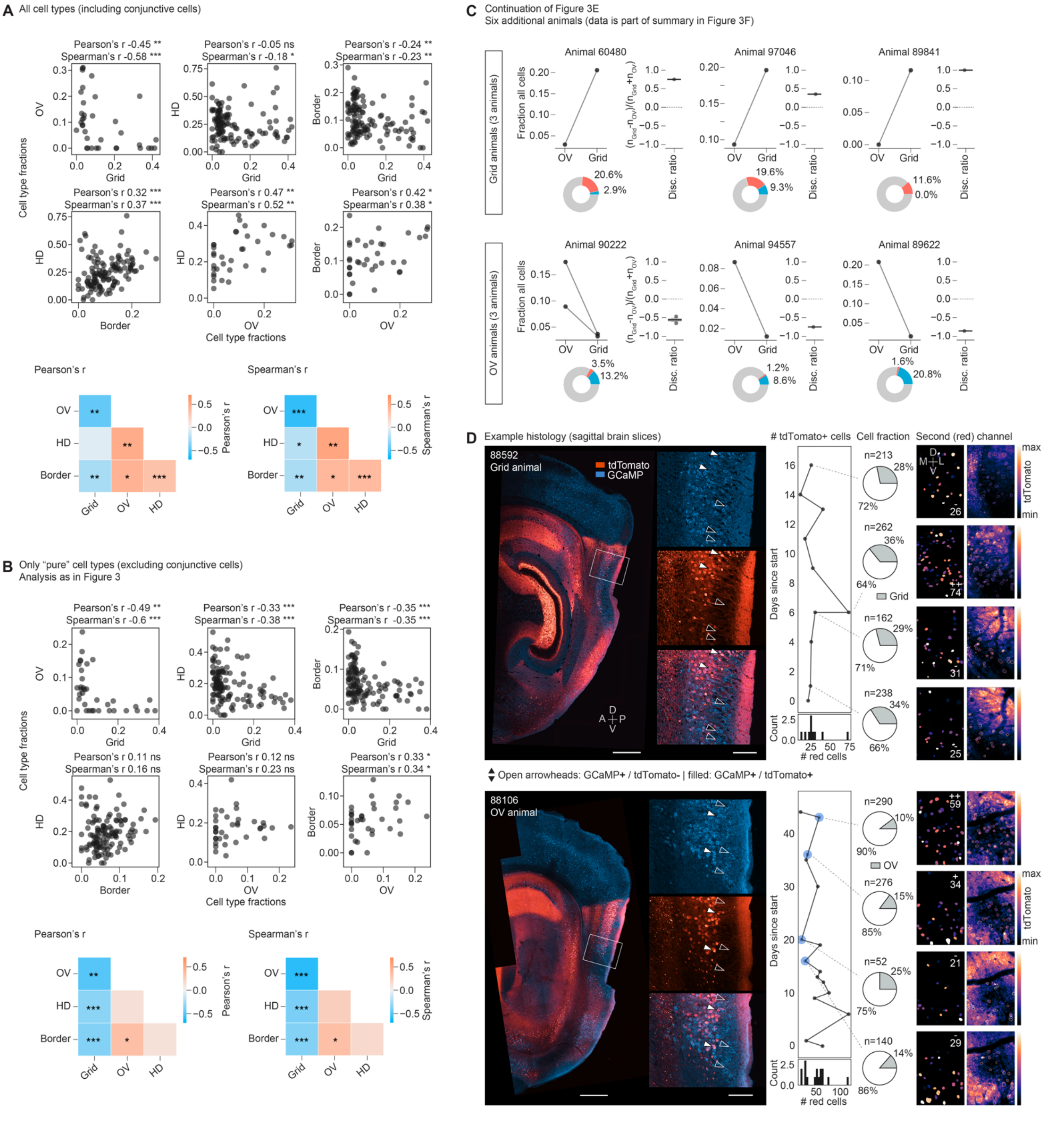


**Figure S3. Correlation of cell type fractions across cell types and quantification of tdTomato positive cells in relation to the fraction of Grid and OV cells in two example animals. Related to Figure 3.**

**(A)** Cell type fractions across all cells per session (comparison as in Figure 3C). Conjunctive cells, i.e., cells that cross the threshold for more than one cell class (95th percentile shuffling cutoff for Grid, HD, Border and combined criteria for OV), are included. Pearson's and Spearman's correlation (r values) and significance tests are indicated in the top right corner of each scatter plot (15 animals, 124 sessions, filtered by MEC, minimum cutoff of 15 cells after filtering). Heatmap on the bottom shows summary of correlation values shown in top graphs. Pearson's correlation: Grid vs. Border: r=-0.24, $n_{Grid}=n_{Border}=124$, p=0.008**; Grid vs. HD: r=-0.05, $n_{Grid}=n_{HD}=124$ p=0.55ns; Grid vs. OV: r=-0.45, $n_{Grid}=n_{OV}=36$, p=0.0054**; OV vs. Border: r=0.42, $n_{OV}=n_{Border}=36$, p=0.011*; OV vs. HD: r=0.47, $n_{OV}=n_{HD}=36$, p=0.0038**; HD vs. Border: r=0.32, $n_{HD}=n_{Border}=124$, p=0.0003***. Spearman's correlation: Grid vs. Border: r=-0.23, $n_{Grid}=n_{Border}=124$, p=0.0093**; Grid vs. HD: r=-0.18, $n_{Grid}=n_{HD}=124$, p=0.041*; Grid vs. OV: r=-0.58, $n_{Grid}=n_{OV}=36$, p=0.0002***; OV vs. Border: r=0.38, $n_{OV}=n_{Border}=36$, p=0.021*; OV vs. HD: r=0.52, $n_{OV}=n_{HD}=36$, p=0.0012**; HD vs. Border: r=0.37, $n_{HD}=n_{Border}=124$, p= 2.37e-05***; ns not significant (p>0.05).

**(B)** Same as in (A) but excluding conjunctive cells (same as in Figure 3C and D). Pearson's correlation: Grid vs. Border: r=-0.35, $n_{Grid}=n_{Border}=124$, p=5.32e-05***; Grid vs. HD: r=-0.33, $n_{Grid}=n_{HD}=124$, p=0.00017***; Grid vs. OV: r=-0.49, $n_{Grid}=n_{OV}=36$, p=0.00240**; OV vs. Border: r=0.33, $n_{OV}=n_{Border}=36$, p=0.048*; OV vs. HD: r=0.12, $n_{OV}=n_{HD}=36$, p=0.49 *ns*; HD vs. Border: r=0.11, $n_{HD}=n_{Border}=124$, p=0.23 *ns*. Spearman's correlation: Grid vs. Border: r=-0.35, $n_{Grid}=n_{Border}=124$, p=5.47e-05***; Grid vs. HD: r=-0.38, $n_{Grid}=n_{HD}=124$ p=1.27e-05***; Grid vs. OV: r=-0.596, $n_{Grid}=n_{OV}=36$, p=0.00013***; OV vs. Border: r=0.34, $n_{OV}=n_{Border}=36$, p=0.046*; OV vs. HD: r=0.23, $n_{OV}=n_{HD}=36$, p=0.17 *ns*; HD vs. Border: r=0.16, $n_{HD}=n_{Border}=124$, p=0.084 *ns*; ns not significant (p>0.05).

**(C)** Visualization as in Figure 3E. Fraction of (pure) grid and OV cells and discrimination index over grid and OV cells in 6 additional animals (included in the summary in Figure 3F), with higher fractions of grid cells than OV cells (top row, "Grid animals") or the opposite (bottom row, "OV animals"). Pie charts underneath each line plot show the average percentage of grid and OV cells across sessions. Boxes in box plots extend from lower to upper quartile values of the data, with a vertical line indicating the population median, whiskers indicate first to 99$^{th}$ percentile. A dashed line is drawn at 0.

**(D)** Left: Confocal images of a sagittal brain slice in two example animals (blue: GCaMP6 (aGFP), red: tdTomato (native fluorescence)). Animal 88592 (top) had multiple grid cells per session ("Grid animal"), while 88106 (bottom) had multiple OV, but hardly any grid cells ("OV animal"). Insets show regions of MEC where the impression of the GRIN+prism implant is visible (approximate location of *in vivo* imaging). Arrow heads in insets point to GCaMP + tdTomato double positive cells (filled arrowheads) and GCaMP only labeled cells (unfilled arrowheads). TdTomato positive cells in these two example animals are enriched in layer III as opposed to layer II. See also Figure S1G for histology in 3 additional animals injected with retro-tdTomato. Counting cells in the secondary (red) imaging channel thereby gives an indication of whether imaging was preferentially targeting layer III or II in each recording. Scale bars: 500 µm, insets: 125 µm.

Middle: Timeline of recorded imaging sessions (days since start of imaging in each animal) and number of recorded red (tdTomato+) cells. The distribution across all recording sessions is shown on the bottom of each timeline. Pie charts towards the right highlight four example sessions for which the percentage of grid (top, 95$^{th}$ shuffling cutoff), and OV (bottom) cells is shown (blue circles indicate object sessions, total number of cells (n) shown on top of each pie chart). No apparent relation between recording date / number of red cells (i.e., cortical layer) and the fraction of grid / OV cells is visible.

Right: Quantification of red cells for the four example sessions indicated in the middle. CellPose was used to extract tdTomato+ cells from bleedthrough corrected average projections of the second (red) channel. Left column: CellPose result (cell masks, number of detected cells is indicated in top right; -: low, +: high, ++: highest). Right column: overlay of cell mask outlines on average projection of second (red) channel, colorbar indicates values in average projection (minimum to maximum).

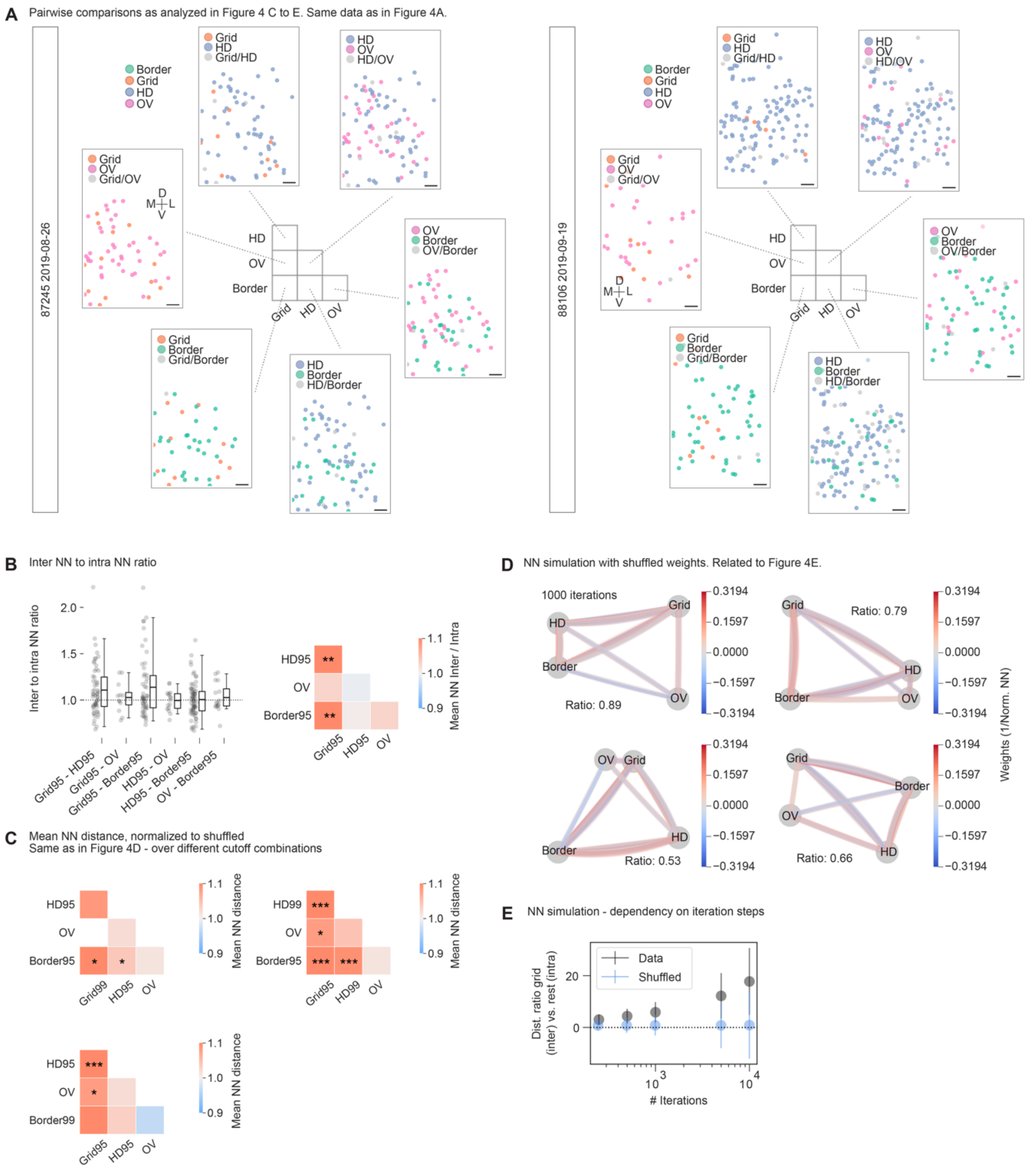


**Figure S4. Extended data for anatomical distribution of functional cell types in MEC. Related to Figure 4.**

**(A)** Example cell maps (same data as shown in Figure 4A) of two recordings in animal 87245 (left, 322 cells) and 88106 (right, 309 cells), showing the distribution of border (green), grid (orange), HD (blue), and OV (pink) cells in pairwise combination (i.e., as analyzed in Figure 4C to E). Cells are color labeled if they crossed shuffling cutoff criteria for one of the two cell types in each comparison and light grey if they met both cutoffs (the latter were excluded from the distance statistics). Tissue orientation is indicated by cross (D-dorsal, L-lateral, V-ventral and M-medial), Scale bars 50 µm.

**(B)** Inter to intra NN ratio measurements across sessions. Session and cell filtering as in Figure 4D. The ratios of average inter-cell class distances (first nearest neighbors, as in Figure 4B to D) over intra-cell class distances (first nearest neighbors) were analyzed in size matched populations. Ratios above one indicate that cells were on average farther apart in between cell classes than within a cell class. The trends are in line with normalized NN distances shown in Figure 4D. Left: Ratios across sessions. Kruskal-Wallis H=5.451, n=6 groups, p=0.36 ns; *Right*: Color coded average of results on the left. Stars indicate results of two-sided one sample t-test against population mean of 1 (*95* indicates the $95^{th}$ percentile shuffling cutoff used as inclusion criterion; p<0.05 *, p<0.01 **, p<0.001 ***; Grid/HD: 12 animals, mean ± SD: 1.11±0.27, t=2.90, $n_{Grid/HD}$=56, p=0.0054**; Grid/OV: 5 animals, mean ± SD: 1.03±0.16, t=0.567, $n_{Grid/OV}$=12, p=0.58 ns; Grid/Border: 12 animals, mean ± SD: 1.14±0.33, t=2.946, $n_{Grid/Border}$=51, p=0.0049**; HD/Border: 13 animals, mean ± SD: 1.01±0.19, t=0.226, $n_{HD/Border}$=68, p=0.82 ns; HD/OV: 6 animals, mean ± SD: 0.99±0.11, t=-0.19, $n_{HD/OV}$=16, p=0.85 ns; OV/Border: 7 animals, mean ± SD: 1.03±0.15 t=0.675, $n_{OV/Border}$=16, p=0.51 ns); ns: not significant (p>0.05).

**(C)** Same as in Figure 4D (right), but over a different combination of cell class cutoff criteria (95: $95^{th}$ percentile shuffling cutoff, 99: $99^{th}$ percentile shuffling cutoff). Stars indicate results of two-sided one sample t-test against population mean of 1 (p<0.05 *, p<0.01 **, p<0.001 ***).

*Top left*: Changed Grid $95^{th}$ to $99^{th}$ percentile shuffling cutoff; Grid/HD: 7 animals, mean ± SD: 1.09±0.20, t=1.900, $n_{Grid99/HD95}$=21, p=0.072 ns; Grid/OV: 1 animal (white square in plot); Grid/Border: 6 animals, mean ± SD: 1.23±0.32, t=2.854, $n_{Grid99/Border95}$=17, p=0.011*; HD/Border: 13 animals, mean ± SD: 1.04±0.15, t=2.260, $n_{HD95/Border95}$=68, p=0.027*; HD/OV: 6 animals, mean ± SD: 1.02±0.07, t=1.065, $n_{HD95/OV}$=16, p=0.30 ns; OV/Border: 7 animals, mean ± SD: 1.01±0.13, t=0.395, $n_{OV/Border95}$=16, p=0.7 ns;

*Top right*: Changed HD $95^{th}$ to $99^{th}$ percentile shuffling cutoff; Grid/HD: 12 animals, mean ± SD: 1.12±0.19, t=4.610, $n_{Grid95/HD99}$=52, p=2.73e-05***; Grid/OV: 5 animals, mean ± SD: 1.08±0.12, t=2.211, $n_{Grid95/OV}$=12, p=0.049*; Grid/Border: 12 animals, mean ± SD: 1.15±0.24, t=4.324, $n_{Grid95/Border95}$=51, p=7.31e-05***; HD/Border: 12 animals, mean ± SD: 1.09±0.19, t=3.671, $n_{HD99/Border95}$=58, p=0.00053***; HD/OV: 6 animals, mean ± SD: 1.04±0.13, t=1.266, $n_{HD99/OV}$=15, p=0.23 ns; OV/Border: 7 animals, mean ± SD: 1.01±0.13, t=0.395, $n_{OV/Border95}$=16, p=0.7 ns;

*Bottom left*: Changed Border $95^{th}$ to $99^{th}$ percentile shuffling cutoff; Grid/HD: 12 animals, mean ± SD: 1.13±0.18, t=5.364, $n_{Grid95/HD95}$=56, p=1.67e-06***; Grid/OV: 5 animals, mean ± SD: 1.08±0.12, t=2.211, $n_{Grid95/OV}$=12, p=0.049*; Grid/Border: 7 animals, mean ± SD: 1.14±0.34, t=1.996, $n_{Grid95/Border99}$=24, p=0.058 ns; HD/Border: 7 animals, mean ± SD: 1.03±0.08, t=1.628, $n_{HD95/Border99}$=19, p=0.12 ns; HD/OV: 6 animals, mean ± SD: 1.02±0.07, t=1.065, $n_{HD95/OV}$=16, p=0.3 ns; OV/Border: 4 animals, mean ± SD: 0.97±0.08, t=-1.088, $n_{OV/Border99}$=8, p=0.31 ns; ns: not significant (p>0.05).

**(D)** Analysis as in Figure 4E. Spring loaded network model with average NN distances acting as weights in between nodes. Four examples with shuffled weights as input are shown, 1000 iterations (simulation steps) each. The inter to intra (Grid vs. Rest) distance ratios are shown for each of the 4 examples.

**(E)** Dependence of the result of the spring-loaded network model on the number of iterations (simulation steps). The inter to intra (Grid vs. Rest) ratio and distribution (Mean ± SD) is shown for data and shuffled (permuted) weights across [250, 500, 1000, 5000, 10000] iterations (simulation steps) (1000 simulations each). Data in Figure 4E and Figure S4D are shown with 1000 iterations and results tend to become more extreme for even more iterations.

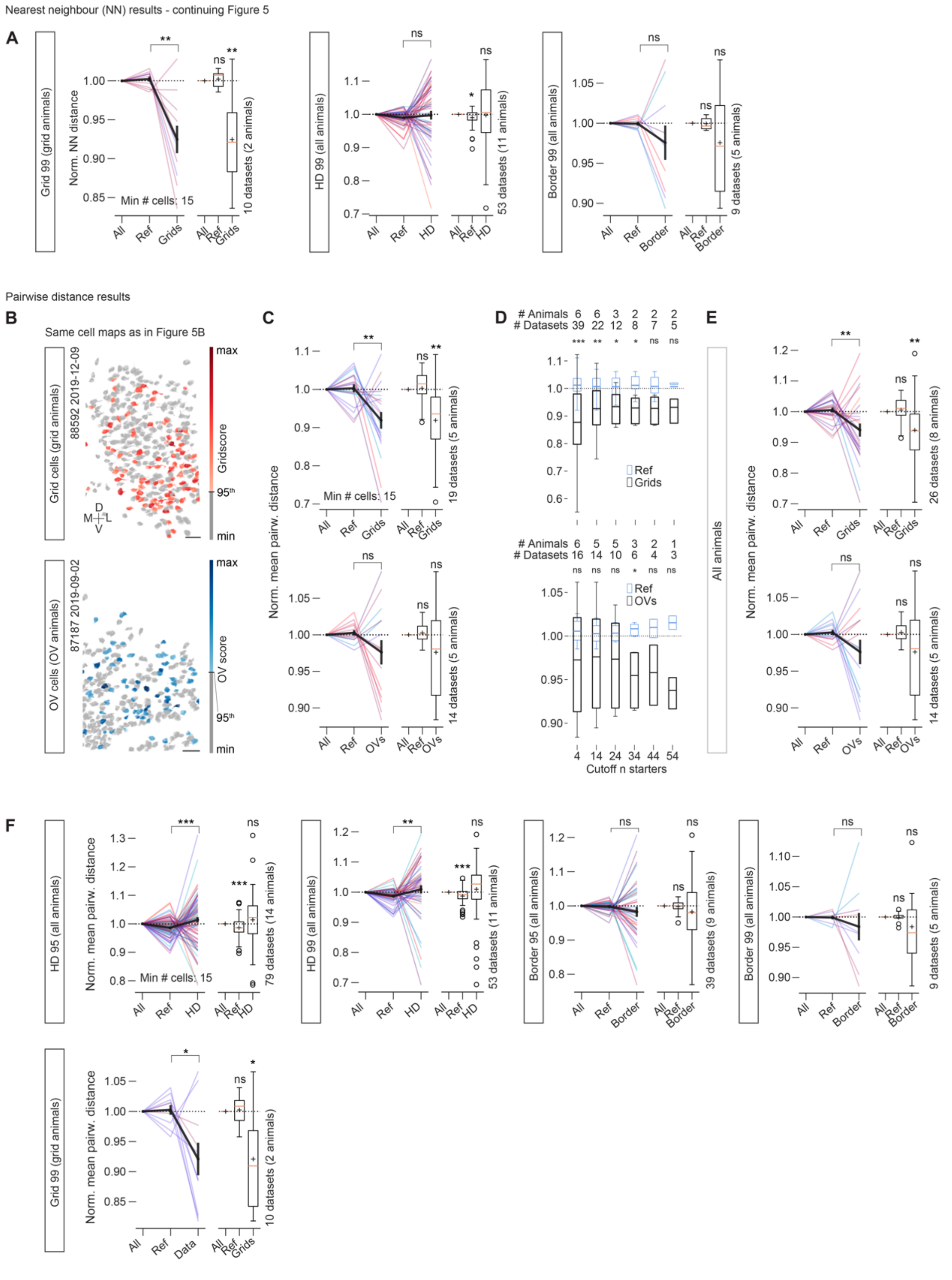


**Figure S5. Extended analysis for clustering of cell classes: NN and pairwise distance analyses. Related to Figure 5.**

(A) nearest neighbor (NN) analyses as in Figure 5 (different cutoffs); (B)-(F): pairwise distance analyses. (B), (C) and (D) show “Grid animals” on top and “OV animals” on bottom.

**(A)** Continuation of nearest neighbor (NN) analyses in Figure 5. As for Figure 5, mean NN distances are normalized to a distribution of cells randomly picked out of all cell locations available for each session and of the same population size as the population of functional cell types under investigation ("All"). A second comparison is shown (reference "Ref"), which consists of cells that were explicitly not part of the cell population under investigation. Left: Grid, cutoff 99$^{th}$ percentile shuffling distribution: *Left:* data over 2 "grid" animals, two-sided Mann-Whitney U=89, $n_{Ref}$=$n_{Grid99}$=10, p=0.004**, Wilcoxon signed-rank test (against 1.): Ref: Z=20, $n_{Ref}$=10, p=0.49 ns Grid99: Z=2 $n_{Grid99}$=10, p=0.0059**; *Middle*: HD, cutoff 99$^{th}$ percentile shuffling distribution: data over 11 animals, Mann-Whitney U=1222, $n_{Ref}$=$n_{HD99}$=53, p=0.25 ns, Wilcoxon signed-rank test (against 1.): Ref: Z=427, $n_{Ref}$=53, p=0.011*, HD99: Z=680 $n_{HD99}$=53, p=0.75 ns; *Right*: Border, cutoff 99$^{th}$ percentile shuffling distribution: data over 5 animals, Mann-Whitney U=54, $n_{Ref}$=$n_{Border99}$=9, p=0.25 ns, Wilcoxon signed-rank test (against 1.): Ref: Z=20, $n_{Ref}$=9, p=0.82 ns, Border99: Z=13, $n_{Border99}$=9, p=0.30 ns; ns: not significant (p>0.05).

**(B)** Two sessions (the same as shown in Figure 5B) in two example animals in which multiple grid cells ("Grid animal", top) or multiple OV cells ("OV animal", bottom) were recorded (color coded by gridscore or OV score). Cells that did not meet cutoff criteria (grid: 95$^{th}$ percentile shuffling) are color coded in grey, animal number and recording date are indicated in top left, anatomical orientation indicated in bottom left (M-medial, L-lateral, V-ventral, D-dorsal). Scale bars 50 µm.

**(C)** Normalized mean pairwise anatomical distance of grid cells in grid animals (top) and OV cells in object vector animals (bottom). The minimum cutoff for number of functional cells (grid cells in grid animals and OV cells in OV animals) is 15. Pairwise distances are normalized to a distribution of cells randomly picked out of all cell locations available for each session and of the same population size as the population of functional cell types under investigation ("All"). A second comparison is shown (reference "Ref"), which consists of cells that were explicitly not part of the cell population under investigation (i.e., non-grid or non-OV cells; same population size as grid / OV cells).

Left: Each line indicates one recording session and colors represent animals. Thick black lines show average and SEM. Grid cells appear to cluster, i.e., pairwise distances are shifted below 1 for grid cells ("Grid" mean ± SEM: 0.92 ± 0.02), while there is a similar trend visible, but no statistical significance, for clustering in OV cells ("OV" mean ± SEM: 0.98 ± 0.02). Box plots show a summary across sessions for data on the left (boxes extend from lower to upper quartile values of the data, with an orange line at the median, whiskers indicate 1$^{st}$ to 99$^{th}$ percentile, outliers are shown as open black circles, black plus signs indicate mean). Vertical labeling towards the right shows the total number of animals and datasets that were used in each comparison. Two-sided Mann-Whitney Grid (top) U=284, $n_{Ref}$=$n_{Grid}$=19, p=0.0026**; Wilcoxon signed-rank test (against 1.): Ref: Z=79, $n_{Ref}$=19, p=0.54 ns; Grid: Z=20, $n_{Grid}$ =19, p=0.0014**; OV (bottom) Two-sided Mann-Whitney U=121, $n_{Ref}$=$n_{OV}$=14, p=0.30 ns; Wilcoxon signed-rank test (against 1.): Ref: Z=44, $n_{Ref}$ =14, p=0.63 ns; OV: Z=33, $n_{OV}$=14, p=0.24 ns; ns: not significant (p>0.05).

**(D)** Comparison as in (C) across different cutoffs (minimum number of functional cells, from 4 to 54) of cells (Grid (top) or OV (bottom)). "Ref" and "Grid"/"OV" summaries are shown as box plots (normalization as in (C)). The number of animals and sessions that remain after filtering are shown above each plot for each comparison. Significance (p < 0.05 *, p < 0.01 **, p < 0.001 ***) indicates results of two-sided Mann-Whitney U test Grid/OV vs. Ref. Boxes extend from lower to upper quartile values of the data, with a thick line at the mean, whiskers indicate 1$^{st}$ to 99$^{th}$ percentile. The observed clustering appears stable over different samples of starter populations in grid cells and trends are visible for OV cells, becoming statistically significant at 34 cells. "Grid animals" mean±SEM for grid cell data and Mann-Whitney U test results: Cutoff 4 cells: 0.88 ± 0.03, U=1225, $n_{Ref}$=$n_{Grid}$=39, p=3.54e-06***; Cutoff 14 cells: 0.93 ± 0.02, U=367, $n_{Ref}$=$n_{Grid}$=22, p=0.0035**; Cutoff 24 cells: 0.93 ± 0.02, U=112, $n_{Ref}$=$n_{Grid}$=12, p=0.023*; Cutoff 34 cells: 0.93 ± 0.02, U=53, $n_{Ref}$=$n_{Grid}$=8, p=0.031*; Cutoff 44 cells: 0.93 ± 0.02, U=39, $n_{Ref}$ =$n_{Grid}$=7, p=0.07 ns; Cutoff 54 cells: 0.93 ± 0.03, U=19, $n_{Ref}$ =$n_{Grid}$=5, p=0.21 ns; "OV animals" mean±SEM for OV cell data and Mann-Whitney U test results: Cutoff 4 cells: 0.97 ± 0.02, U=160, $n_{Ref}$=$n_{OV}$=16, p=0.24 ns; Cutoff 14 cells: 0.98 ± 0.02, U=121, $n_{Ref}$=$n_{OV}$=14, p=0.30 ns; Cutoff 24 cells: 0.97 ± 0.02, U=64, $n_{Ref}$=$n_{OV}$=10, p=0.31 ns; Cutoff 34 cells: 0.95 ± 0.02, U=31, $n_{Ref}$=$n_{OV}$=6, p=0.045*; Cutoff 44 cells: 0.96 ± 0.02, U=13, $n_{Ref}$=$n_{OV}$=4, p=0.19 ns; Cutoff 54 cells: 0.94 ± 0.02, U=9, $n_{Ref}$=$n_{OV}$=3, p=0.08 ns; ns: not significant (p>0.05).

**(E)** Comparison as in (C) but over all animals and recordings (no preselection for animals, which fall into the categories grid or OV animals). Grid mean ± SEM: 0.94 ± 0.02, Two-sided Mann-Whitney Grid vs. Ref U=496, $n_{Ref}$=$n_{Grid}$=26, p=0.004**; Wilcoxon signed-rank test (against 1.): Ref: Z=137, $n_{Ref}$=26, p=0.33 ns; Grid: Z=74, $n_{Grid}$=26, p=0.00994**. OV mean ± SEM: 0.98 ± 0.02, Two-sided Mann-Whitney OV vs. Ref U=121, $n_{Ref}$=$n_{OV}$=14, p=0.30 ns; Wilcoxon signed-rank test (against 1.): Ref: Z=44, $n_{Ref}$=14, p=0.63 ns; OV: Z=33, $n_{OV}$=14, p=0.24 ns. Vertical labeling towards the right shows the total number of animals and datasets that were used in each comparison; ns: not significant (p>0.05).

**(F)** As in (C) and (E) but for head direction (HD) and border cells ($95^{th}$ or $99^{th}$ percentile shuffling cutoffs). Compared to grid cells no clean trends are observable for these cell classes.

HD95 mean ± SEM: 1.01 ± 0.01, Two-sided Mann-Whitney HD95 vs. Ref U=2153, $n_{Ref}$=$n_{HD95}$=79, p=0.00077***; Wilcoxon signed-rank test (against 1.): Ref: Z=873, $n_{Ref}$=79, p=0.00055***; HD95: Z=1225, $n_{HD}$=79, p=0.083 ns; HD99 mean ± SEM: 1.01 ± 0.01, Two-sided Mann-Whitney HD99 vs. Ref U=886, $n_{Ref}$=$n_{HD99}$=53, p=0.0011**; Wilcoxon signed-rank test (against 1.): Ref: Z=337, $n_{Ref}$=53, p=0.00081***; HD: Z=507, $n_{HD99}$=53, p=0.065 ns; Border95 mean ± SEM: 0.98 ± 0.01, Two-sided Mann-Whitney Border95 vs. Ref U=900, $n_{Ref}$=$n_{Border95}$=39, p=0.16 ns; Wilcoxon signed-rank test (against 1.): Ref: Z=375, $n_{Ref}$=39, p=0.83 ns; Border95: Z=287, $n_{Border95}$=39, p=0.15 ns; Border99 mean ± SEM: 0.98 ± 0.02, Two-sided Mann-Whitney Border99 vs. Ref U=48, $n_{Ref}$=$n_{Border99}$=9, p=0.54 ns; Wilcoxon signed-rank test (against 1.): Ref: Z=20, $n_{Ref}$=9, p=0.82 ns; Border99: Z=17, $n_{Border99}$=9, p=0.57 ns;

Last row: Grid cells > $99^{th}$ percentile cutoff in grid animals. Data over 2 “grid” animals, Mann-Whitney U test Grid vs. Ref: U=78, $n_{Ref}$=$n_{Grid99}$=10, p=0.038*; Wilcoxon signed-rank test (against 1.): Ref: Z=24, $n_{Ref}$ =10, p=0.77 ns, Grid: Z=6, $n_{Grid99}$=10, p=0.027*; ns: not significant (p>0.05).

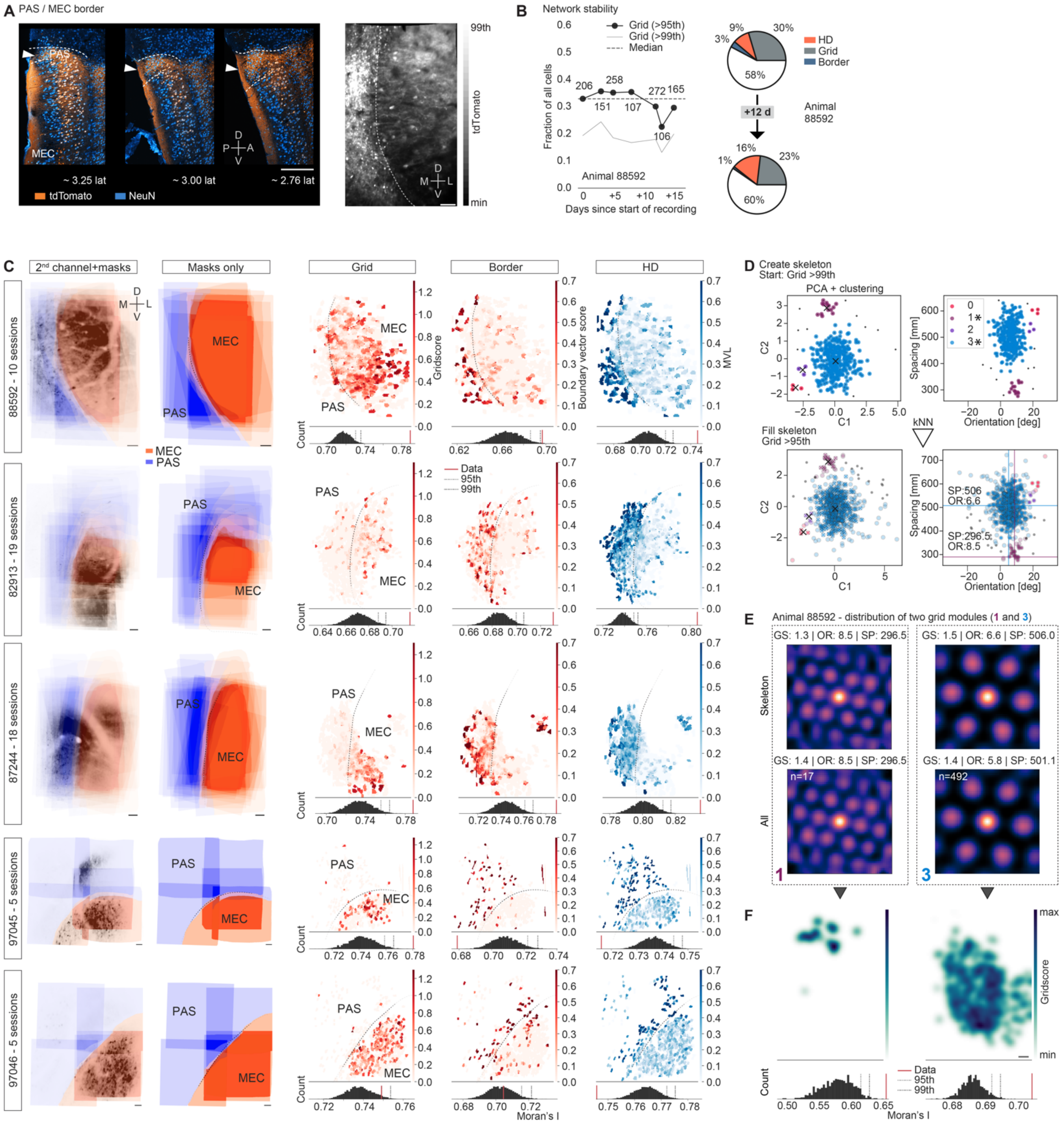


**Figure S6. Stability of functional network composition and identification of PAS/MEC boundary in histology, *in vivo* topographic tuning maps, and example of grid module distribution. Related to Figure 6.**

**(A)** Left: Confocal images (maximum intensity projections) of immunostaining in three sagittal brain slices from one animal (cyan - aNeuN, orange - retroAAV tdTomato). Approximate mediolateral position (distance from midline) indicated at the bottom ("lat", unit is mm). Dashed lines indicate the boundary between parasubiculum (PAS) and overlying cortex (top) and the boundary between PAS and medial entorhinal cortex (MEC) (bottom). The boundaries were distinguishable by changes in the pattern of tdTomato labeled neuropil and cellular labeling in MEC. Arrowhead points towards the expanding territory of PAS (lateral to medial); P - posterior, A - anterior, D - dorsal, V - ventral, scale bar 250 μm. Right: Example *in vivo* imaging field of view (secondary, “red” channel) in the same animal as shown on the left. The boundary between MEC and adjacent PAS is indicated by a dashed line); M - medial, L - lateral, D - dorsal, V - ventral, scale bar 50 μm; color bar shows range of values (minimum to maximum) in second channel projection.

**(B)** The composition of functional cell types stayed relatively stable over multiple recorded sessions, here shown for one example animal in which multiple grid cells were co-recorded over many days. Left: Fraction of grid cells meeting either 95$^{th}$ (black line and dots) or 99$^{th}$ percentile (grey line) shuffling cutoffs (dashed line indicates median fraction of cells meeting the 95$^{th}$ percentile cutoff over all sessions). Numbers on top indicate the total number of recorded cells (filtered by SNR). Right: Pie charts showing the fraction of functional cell types in two example sessions of the same animal as on the left. The fractions stay relatively stable.

**(C)** For every recording, anatomical masks were drawn manually while visualizing the second (red) channel average projection. After stitching recordings (composite FOV based on user defined anatomical landmarks), these masks align to form consistent boundaries across PAS and MEC. Left to right: average projection of stitched and aligned second channel images (as in Figure 6A, right) with anatomical masks overlaid; masks only (anatomical masks are shown with transparency such that the intensity of the color indicates the number of total sessions that were stacked on top of each other); topographic tuning maps as in Figure 6B, dashed lines indicate approximate location of PAS / MEC (derived from annotations shown on the left. Scale bars 50 μm.

**(D)** Grid module extraction. Example data from one animal with many grid cells. After dimensionality reduction (PCA, left, C1 and C2 - principal components 1 and 2), unsupervised clustering is run on a core set of “pure” grid cells (>99$^{th}$ percentile cutoff) to extract seed clusters (“skeleton”, top). A k-nearest neighbor classifier (kNN) is trained on this data; this classifier is subsequently used to expand each module by adding grid cells with >95$^{th}$ (and <99$^{th}$) percentile shuffling cutoff (bottom). Four candidate modules were extracted / expanded for this example animal. The two main modules (those with high numbers of classified cells) – modules 1 (spacing ~ 297 mm and orientation ~ 9 degrees, magenta dots, lines indicate means) and 3 (spacing ~ 506 mm and orientation ~ 7 degrees, cyan dots, lines indicate means) – are shown in (E).

**(E)** Average spatial autocorrelation maps of modules 1 and 3 (extracted in (D)). Top: Module properties of a subset of “pure” grid cells (> 99$^{th}$ percentile shuffling cutoff) used as “Skeleton” and all cells on bottom (“All”) (GS - Grid Score, OR - orientation [degrees], SP - spacing [mm], total number of cells (n=17 for smaller (left, module 1) and n=492 for larger (right, module 3) module)).

**(F)** Modules extracted in (D) and (E), mapped over anatomical space (Left: Module 1, Right: Module 3 over the same FOVs). Topographical tuning maps, Moran’s I distributions and cutoffs (bottom) as in Figure 6B and Figure S6C. Colorbar shows grid tuning mapped from minimum to maximum for the scoremap average projection (7 sessions). Scale bar: 50 μm.
