## Supplementary material for "Functional network topography of the medial entorhinal cortex": Key Resources Table

**This file includes:**

Key Resources Table

### KEY RESOURCES TABLE

| REAGENT or RESOURCE | SOURCE | IDENTIFIER |
|---|---|---|
| **Antibodies** | | |
| Mouse monoclonal anti-NeuN, clone A60 (primary antibody) | Merck | Cat# MAB377, RRID:AB_2298772 |
| Rabbit polyclonal anti-GFP (primary antibody) | Thermo Fisher Scientific | Cat# A-11122, RRID:AB_221569 |
| Rabbit anti-mouse IgG (H+L), Alexa Fluor 647 (secondary antibody) | Abcam | Cat# ab169348, RRID:AB_2893161 |
| Donkey anti-rabbit IgG (H+L), Alexa Fluor 488 (secondary antibody) | Thermo Fisher Scientific | Cat# A-21206, RRID:AB_2535792 |
| **Bacterial and Virus Strains** | | |
| AAV1.Syn.GCaMP6m | UPenn Vector Core | Cat#AV-1-PV2823 |
| AAVretro.CAG.tdTomato | Addgene, (Tervo et al., 2016) | RRID:Addgene_59462 |
| **Experimental Models: Organisms/Strains** | | |
| Mouse wild type C57BL/6JBomTac | Taconic | Cat#B6JBOM; RRID: IMSR_TAC:b6jbom |
| Mouse transgenic B6;DBA-Tg(tetO-GCaMP6s)2Niell/J | Wekselblatt et al., 2016 | RRID:IMSR_JAX:024742 |
| Mouse transgenic B6.Cg-Tg(Camk2a-tTA)1Mmay/DboJ | Wekselblatt et al., 2016 | RRID:IMSR_JAX:007004 |
| **Software and Algorithms** | | |
| Python | Van Rossum and Drake, 2009 | https://www.python.org/ |
| Napari | Sofroniew et al., 2021 | doi:10.5281/zenodo.3555620 |
| Fiji (ImageJ) | Schindelin et al., 2012 | http://fiji.sc; RRID: SCR_002285 |
| ScanImage 5.5 | Vidrio Technologies | http://scanimage.vidriotechnologies.com/display/SI5/ScanImage+5 |
| Suite2p python | Pachitariu et al., 2017 | https://github.com/MouseLand/suite2p |
| SEP: Python library for Source Extraction and Photometry | Barbary, 2016 | https://github.com/kbarbary/sep |
| CircStat | Berens, 2009 | https://github.com/circstat/pycircstat |
| PySAL: Spatial Analysis Library for open source, | Rey and Anselin, 2010 | https://pysal.org/ |

| | | |
|---|---|---|
| cross platform, Geospatial Data Science | | |
| Scikit-learn | Pedregosa et al., 2011 | https://scikit-learn.org/ |
| CMasher: Scientific colormaps for making accessible, informative and "cmashing" plots | Van der Velden, 2020 | https://github.com/1313e/CMasher |
| SciPy | Virtanen et al., 2020 | https://www.scipy.org/ |
| Datajoint | Yatsenko et al., 2015 | https://github.com/datajoint/datajoint-python |
| Brainreg-segment | Tyson et al., 2021 | https://github.com/brainglobe/brainreg-segment |
